# Mucosal-associated invariant T cells support IL-15-dependent Treg response in skin injury and promote resolution of skin inflammation

**DOI:** 10.64898/2026.06.28.735088

**Authors:** Grace E. Crossland, Andre Armero, Vianey Chavez, Zachary T. Peters, Lennard Ostendorf, Shriya Kannan, Lindsay K. Mendyka, Olivia F. Brooker, Kaitlyn Dowling, Himanshu B. Goswami, Dorothea Barton, Sonia Leach, Fred W. Kolling, Pamela C. Rosato, Mark S. Sundrud, Theresa T. Lu, Deepak A. Rao, Michael G. Constantinides, Sladjana Skopelja-Gardner

## Abstract

Mucosal-associated invariant T (MAIT) cells are enriched at barrier sites, but their role in autoimmune skin inflammation remains unknown. Using cutaneous lupus as a model, we identify MAIT cells as protective regulators of skin inflammation and as critical upstream modulators of regulatory T cells (Treg). Topical MAIT cell activation with 5-OP-RU induced durable resolution of spontaneous skin lesions in MRL/lpr mice and suppressed TLR7–driven skin inflammation. MAIT cell activation selectively expanded and activated Treg populations in both healthy and lupus-like skin, while suppressing effector T cell cytokine production and cytotoxic programs. This MAIT-Treg axis was also activated in UV light-driven barrier injury in healthy murine and human skin, where MAIT cells were required for UV-elicited Treg expansion and function. In lupus-like skin, local MAIT cell activation restored the defective UVB-induced Treg response and limited CD8+ T cell expansion. Mechanistically, CCR2+ monocyte-derived antigen-presenting cells and IL-15 signaling were required for MAIT cell-driven Treg accumulation and therapeutic benefits of MAIT cells in inflamed skin. These studies identify a MAIT-IL-15-Treg axis that links barrier injury sensing to immune regulation, which is disrupted in cutaneous lupus, and nominate therapeutic MAIT cell activation as an unappreciated strategy for restoring immune homeostasis in inflamed skin.

## Introduction

Mucosal-associated invariant T (MAIT) cells are innate-like lymphocytes enriched in barrier and mucosal tissues and are highly conserved throughout mammalian evolution (*1, 2*). They are defined by a semi-invariant TCR, restricting the repertoire of antigens they recognize to those presented by the MHC-I-related molecule MR1, which is critical to their development in the thymus (*1, 3, 4*). The best-studied MR1 ligands are 5-(2-oxopropylideneamino)−6-D-ribitylaminouracil (5-OP-RU) and 5-(2-oxoethylideneamino)−6-D-ribitylaminouracil (5-OE-RU), metabolites of bacterial riboflavin synthesis (*5, 6*), though endogenous MR1 ligands have been reported (*7*). Due to their innate-like features, MAIT cells can be activated independently of TCR by inflammatory cytokines, including IL-12, IL-18, and type I interferons (IFN-I) (*8–10*).

MAIT cells rapidly sense perturbations in barrier tissues and exhibit diverse effector functions (*11*). They mediate antimicrobial and antiviral responses by producing inflammatory cytokines, including IFN-γ, TNF-α, and IL-17, as well as cytotoxic molecules such as granzymes and perforin (*8, 12–15*). More recently, MAIT cells have been implicated in maintaining barrier tissue integrity and promoting wound repair by supporting stromal cells via TGFβ and amphiregulin (*16–19*). The same properties that enable MAIT cells to respond to diverse infectious and sterile stimuli, including their cytokine-responsiveness, tissue localization, and capacity to promote inflammation or repair, also position them to respond to immune dysregulation. Since tissue dysbiosis, defects in barrier integrity, and an altered cytokine milieu are hallmarks of many autoimmune diseases, MAIT cells are likely to contribute in these settings. Consistent with this notion, MAIT cell frequency is often reduced in the circulation of patients with autoimmune diseases (*20–25*), and some studies report increased numbers in affected tissues (*26–28*). However, whether MAIT cells drive pathology or instead protect tissues, particularly the skin, in inflammatory and autoimmune disease remains unclear.

Therefore, we turned to cutaneous lupus erythematosus (CLE) as a model of chronic autoimmune skin inflammation to understand MAIT cells in skin autoimmunity. Peripheral MAIT cells are reduced but activated in lupus, yet single-cell RNA sequencing studies of CLE skin have failed to capture MAIT cells, limiting assessment of their function in diseased tissue (*24, 25, 29–31*). CLE skin exhibits features that can activate MAIT cells, including elevated IFN-I, microbial dysbiosis, spontaneous lesion formation, and heightened susceptibility to ultraviolet (UV)-induced tissue injury (photosensitivity) (*32–34*). However, MAIT cell biology in CLE remains undefined.

Here, we asked whether MAIT cell activation in CLE-like skin could resolve or prevent disease and sought to define the underlying mechanisms. Using the cognate antigen 5-OP-RU, we found that TCR-driven MAIT cell activation resolved both chronic and acute lupus-like skin lesions *in vivo*. Local MAIT cell stimulation induced cutaneous expansion and activation of immunosuppressive regulatory T cells (Treg), a response lost with genetic deletion of MAIT cells. Skin exposure to a disease-relevant stimulus, UV light, similarly induced MAIT cell expansion and Treg activation, in a MAIT cell-dependent fashion. In CLE-like skin, restoring MAIT cell levels suppressed inflammatory mediators and prevented UV-induced skin flares. Furthermore, antigen-mediated activation of MAIT cells potentiated the UV-induced Treg response in human skin explants. MAIT-cell-driven Treg expansion required IL-15 signaling and CCR2+ monocyte-derived antigen-presenting cells (moAPCs). Together, these findings define a cutaneous MAIT-Treg axis mediated by moAPCs and IL-15, highlighting a potential therapeutic pathway in inflammatory skin disease.

## Results

### Local activation of MAIT cells restores immune regulation to heal active inflammatory skin lesions

In health, MAIT cells support barrier tissue integrity and wound repair (*16–19, 35, 36*). However, they are functionally heterogeneous and can also promote inflammation (*11*). Across autoimmune diseases, MAIT cells are reduced in patients’ peripheral blood (*20–25, 37–39*), prompting the hypothesis that they traffic into tissues and contribute to pathology. Yet, in most autoimmune conditions, particularly those of the skin, MAIT cells have been difficult to identify and study within the affected tissue, leaving it unclear whether they promote disease or protect tissue. To assess whether MAIT cells are protective or pathogenic in inflammatory skin disease, we used MRL/lpr mice, which harbor a *Fas* mutation that drives rapid spontaneous systemic autoimmunity, and allowed them to develop spontaneous skin lesions scored for erythema, scaling, thickness, alopecia, and the extent of skin area involved (*40*). When lesions reached a score >10, we initiated antigen-mediated cutaneous expansion of MAIT cells. In Cohort 1, mice received topical 5-OP-RU or vehicle every 48 hours, together with systemic isotype control or anti-MR1 IgG to block MR1 antigen presentation (**Fig. 1A**). In Cohort 2, mice received topical 5-OP-RU or vehicle every 48 hours for nine total doses and were followed to assess the durability of the response (**Fig. 1A**). In Cohort 1, 5-OP-RU led to near-complete resolution of skin lesions, whereas vehicle-treated mice showed no improvement; MR1 blockade abrogated the therapeutic effect of 5-OP-RU (**Fig. 1B–C**). Lesion resolution following local MAIT cell expansion was sustained for up to 40 days after 5-OP-RU cessation, whereas vehicle-treated lesions failed to resolve and continued to flare (**Fig. 1D**). Back and ear skin epidermal and dermal thickness were reduced in 5-OP-RU + isotype–treated mice relative to mice receiving 5-OP-RU + anti-MR1 IgG (**Fig. 1E**, fig. S1A).

**Figure 1.**
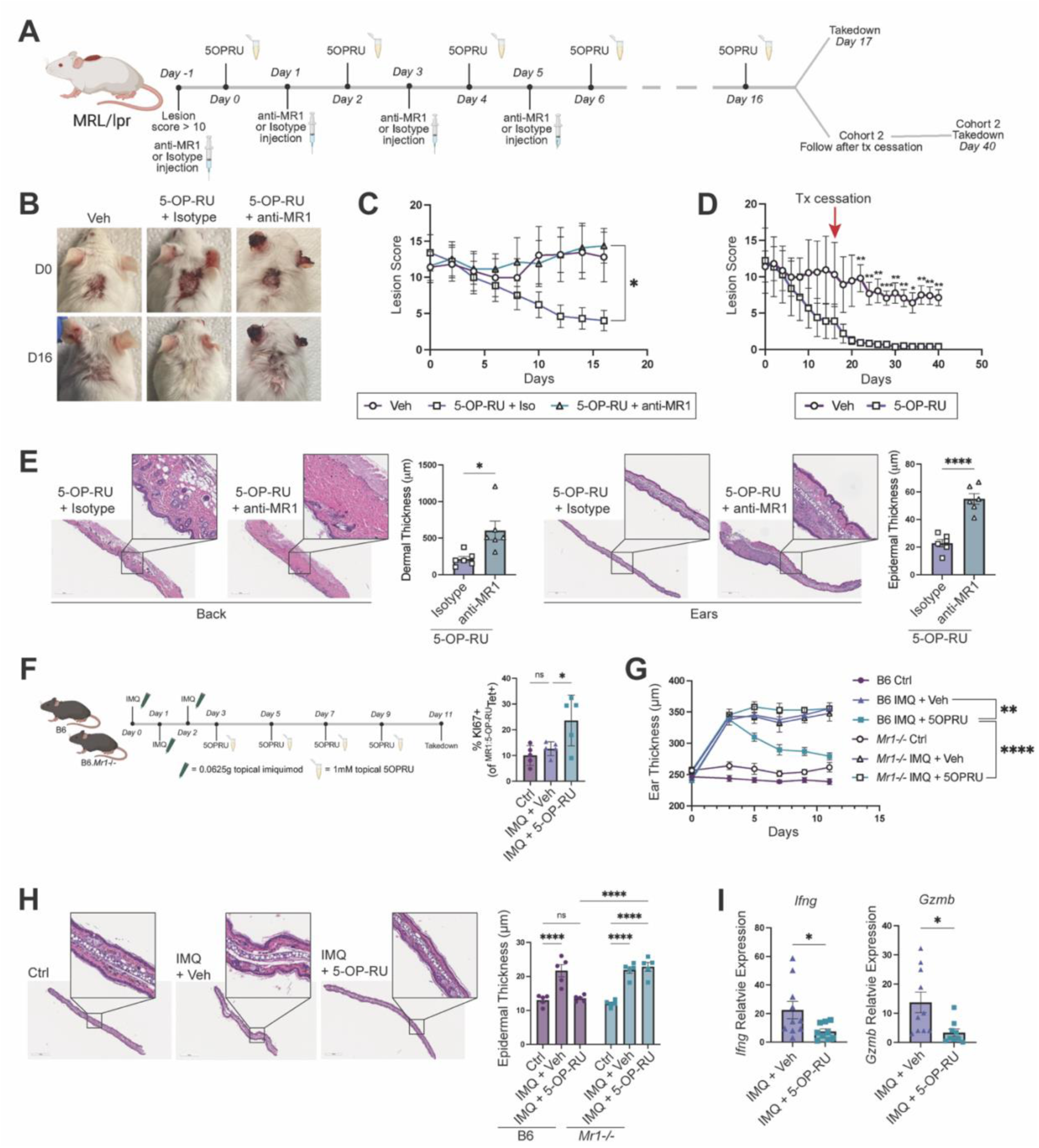
Topical MAIT cell expansion heals active chronic and inflammatory skin lesions. (**A**) Schematic of topical MAIT cell activation in Cohort 1 (B-C) and Cohort 2 (D) of MRL/lpr mice (male and female, 12-18wks). (**B**) Representative images of MRL/lpr skin lesions at treatment initiation (D0) and cessation (D16). (**C**) Lesion scores in Cohort 1 calculated on the degree of erythema, scaling, thickness, and alopecia (n=6/group). (**D**) Lesion scores in Cohort 2 mice treated with 5-OP-RU vs. vehicle over time (n=7/group). (**E**) Representative H&E and corresponding back and ear skin thickness quantification in mice treated with 5-OP-RU with (anti-MR1) or without (Isotype) blockade of antigen presentation. (**F**) Schematic of acute induction of inflammation via topical imiquimod (IMQ) prior to treatment with topical 5-OP-RU or vehicle in B6 and *Mr1-/-* mice and quantification of MAIT cell KI67 expression. (**G**) Ear thickness measurements over the course of treatment (n=5/group). (**H**) Representative H&E and corresponding ear thickness quantification of control or IMQ-treated ears with (5-OP-RU) and without (Veh) MAIT cell expansion. (**I**) Expression of cytotoxic (*Ifng, Gzmb*) genes determined by qPCR relative to *Gapdh* in B6 IMQ-treated ears with (5-OP-RU) or without (Veh) MAIT cell expansion (n=10/group). Data represented as means +/- SEM. Data analyzed with (C, F) one-way ANOVA, (D, I) Welch’s *t-*test, (G, H) two-way ANOVA. *p<0.05, **p<0.01, ***p<0.001, ****p<0.0001. ns, not significant.

Because spontaneous MRL/lpr skin lesions are chronic, with signatures of tissue remodeling and wound healing (*41*), we next asked whether activating MAIT cells could also suppress acute lupus-like skin inflammation. Since high IFN-I levels are a hallmark of lupus and TLR7 is linked to disease etiology (*42, 43*), we employed an acute TLR7 agonist model (imiquimod, IMQ) to mimic a lupus skin flare. IMQ was applied for three consecutive days to induce acute ear inflammation, i.e., an increase in ear thickness (*44*). MAIT cells were activated in the inflamed skin by topical 5-OP-RU, and the change in ear thickness was compared to vehicle-treated ears in both B6 and MAIT cell-deficient *Mr1*-/- mice (**Fig. 1F**, fig. S1B). Strikingly, 5-OP-RU reduced ear thickness to basal levels in B6 skin (**Fig. 1G**). On the contrary, *Mr1*-/- ears remained inflamed regardless of treatment, demonstrating an MR1-MAIT requirement for the anti-inflammatory effects of 5-OP-RU (**Fig. 1G**). Histopathologic analysis showed that 5-OP-RU ameliorated epidermal thickening in B6 but not *Mr1-/-* mice, and decreased expression of cytotoxic mediators, including *Ifng* and *Gzmb* (**Fig. 1H-I**, fig. S1C).

### TCR-driven MAIT cell expansion supports skin Treg and suppresses immune activation

To understand the mechanism by which TCR-mediated MAIT cell activation drives resolution of inflammatory skin lesions, we profiled the skin T cell compartment after 5-OP-RU administration in B6 and *Mr1-/-* skin (**Fig. 2A-B**, fig. S2A, S3A). Topical 5-OP-RU induced ∼3-fold expansion of MAIT cells in the skin (**Fig. 2B-C**). Unexpectedly, we observed a concordant ∼4-fold increase in cutaneous Treg, with no detectable effects on other T cell subsets (**Fig. 2B-D**, fig. S3A-E). Importantly, Treg did not expand in 5-OP-RU-treated *Mr1*-/- skin (**Fig. 2D**). Transcriptomic analysis of FACS-sorted GFP+ Treg from vehicle- or 5-OP-RU-treated skin revealed that antigen-mediated MAIT cell activation induced Treg expansion across different subsets (fig. S3F). Moreover, Treg in 5-OP-RU-treated skin exhibited high expression of genes involved in regulatory functions (e.g., *Ctla4*, *Tigit, Il10, Il2ra, Entpd1*), resulting in a higher overall immunoregulatory gene score (**Fig. 2E**, **Table 1**). Dimensionality reduction of Treg flow cytometry data confirmed that 5-OP-RU stimulated broad expansion across Treg subsets (defined as in (*45*)) and drove MAIT cell-dependent Treg activation, indicated by increased ICOS on Treg from 5-OP-RU-treated B6, but not *Mr1*-/-, skin (**Fig. 2F-G**, fig. S3G).

**Figure 2.**
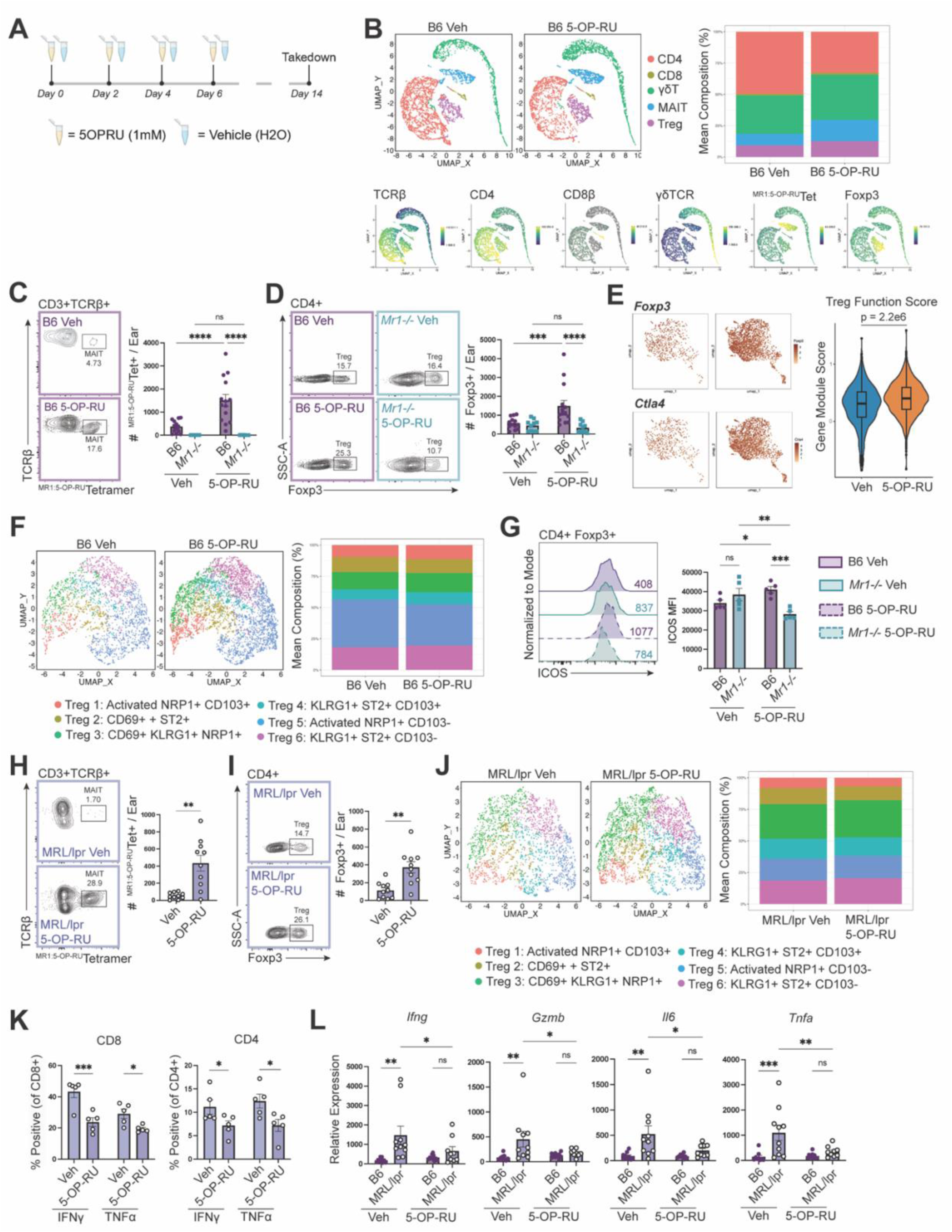
MAIT cell activation supports Treg expansion and suppresses cutaneous inflammation. (**A**) Schematic of MAIT cell expansion via topical 5-OP-RU in B6, *Mr1-/-*, or MRL/lpr mice (female, 8wks). (**B**) Flow cytometry UMAP and corresponding stacked bar plot of T cells from 5-OP-RU or Veh-treated B6 and *Mr1-/-* skin (n=5/group). (**C-D**) Flow cytometry contour plots and quantification of (C) MAIT cells (live CD45+ γδTCR- CD3+ TCRβ+ ^MR1:5-OP-RU^Tetramer+) and (D) Treg (live CD45+ γδTCR- CD3+ TCRβ+ CD4+ Foxp3+) in B6 and *Mr1-/-* skin treated with topical 5-OP-RU or vehicle (n=14/group). (**E**) ScRNAseq feature plots of *Foxp3* and *Ctla4* and transcriptomic function score in Treg from 5-OP-RU- or vehicle-treated B6.*Foxp3-GFP* skin. (**F**) Flow cytometry UMAP and corresponding stacked bar plot of Treg from 5-OP-RU or Veh-treated B6 and *Mr1-/-* skin (n=5/group). (**G**) Flow cytometry histograms and corresponding quantification of ICOS on Treg from 5-OP-RU or Veh-treated B6 and *Mr1-/-* skin (n=5/group). (**H-I**) Flow cytometry contour plots and corresponding quantification of (H) MAIT cells and (I) Treg in 5-OP-RU or Veh-treated MRL/lpr skin (n=10/group). (**J**) Flow cytometry UMAP and corresponding stacked bar plot of Treg from 5-OP-RU or Veh-treated MRL/lpr skin (n=5/group). (**K**) Quantification of CD8+ and CD4+ T cell cytokine production in 5-OP-RU or Veh-treated MRL/lpr skin determined by flow cytometry (n=5/group). (**L**) Expression of cytotoxic (*Ifng, Gzmb*) and inflammatory (*Il6, Tnf*) genes determined by qPCR relative to *Gapdh* in 5-OP-RU or Veh-treated B6 and MRL/lpr skin (n=10/group). Data represented as means +/- SEM. Data analyzed with (C-D, G, L) two-way ANOVA, (E) Wilcoxon rank sum test, or (H-I, K) Welch’s *t-*test. *p<0.05, **p<0.01, ***p<0.001, ****p<0.0001. ns, not significant.

Even in the absence of overt lesions, MRL/lpr skin exhibits heightened immune activation and Treg dysfunction, evidenced by increased frequencies of IFN-γ-producing CD8+ and CD4+ T cells, reduced Treg IL-10 and amphiregulin production, and decreased MAIT cell frequencies (fig. S4A-G). Therefore, we next tested whether TCR-dependent MAIT cell activation in non-lesional MRL/lpr skin could suppress immune activation by restoring regulatory pathways, as seen in healthy B6 skin. Topical 5-OP-RU robustly expanded MAIT cells and induced ∼4-fold increase in Treg numbers (**Fig. 2H-I**). As in B6 skin, Treg expanded across subsets in non-lesional CLE-like skin (**Fig. 2J**). Consistent with an immunoregulatory effect, 5-OP-RU reduced the frequencies of IFN-γ- and TNF-α-producing CD8+ and CD4+ T cells (**Fig. 2K**, fig. S4H-I). It also reduced the expression of cytotoxic (*Gzmb, Ifng*) and inflammatory (*Tnfa, Il6*) genes to levels approaching those in healthy B6 skin (**Fig. 2L**). Together, these findings demonstrate that antigen-driven MAIT cell expansion promotes Treg accumulation even in an inflammatory skin environment and dampens basal immune activation in CLE-like skin.

### MAIT cells are required for UV-induced Treg activation and immune suppression in the skin

Having established a functional MAIT-Treg axis in the context of antigen-driven MAIT cell activation, we next asked whether MAIT cells are required for Treg expansion in response to UVB light, a known stimulus of Treg-dependent immunosuppression in healthy murine skin (*46–48*). Using *ex vivo* human skin explants, we confirmed that a single UVB exposure is likewise sufficient to stimulate local Treg expansion and proliferation in human skin (**Fig. 3A-C**, fig. S5A). Because viable MAIT cells were difficult to recover from human skin, we could not determine whether UVB induced MAIT cell expansion *in situ*. Therefore, we developed a Transwell system in which healthy human skin biopsies were exposed to a single dose of UVB before co-culture with healthy PBMCs (**Fig. 3D**). In this model, UVB-exposed skin induced expansion of MAIT cells and proliferation of both MAIT cells and Treg by 72 hours post-UVB (**Fig. 3E-F**, fig. S5B-D). We therefore hypothesized that, as with antigen-driven MAIT cell activation (**Fig. 2A-C**), MAIT cells are required for UVB-triggered immunoregulatory responses in healthy skin, including Treg expansion. To test this, we exposed B6 and *Mr1-/-* mice to chronic, low-dose UVB and quantified cutaneous MAIT cells and Treg by flow cytometry (**Fig. 3G**, fig. S6A-B). Confirming the human *ex vivo* observations, UVB induced robust MAIT cell expansion in B6 skin *in vivo* (fig. S6A). Strikingly, UVB failed to elicit Treg expansion or proliferation in *Mr1*-/- skin (**Figure 3H-I**, fig. S6B). Treg did not preferentially accumulate in the skin-draining lymph nodes or spleen of *Mr1-/-* mice (fig. S6C-D), suggesting the lack of expansion in the skin is not due to impaired trafficking. Spectral cytometry profiling of Treg after UVB exposure showed expansion of the same six Treg subsets identified following topical 5-OP-RU (**Fig. 3J**, fig. S6E). While UVB induced a proportional increase across Treg subsets in B6 skin, the Treg composition shifted slightly in favor of CD103-negative clusters 5 and 6 in *Mr1*-/- skin, despite no increase in the overall Treg abundance (**Fig. 3J**). Additionally, after UVB exposure, Treg in B6, but not *Mr1-/-* skin, upregulated the effector Treg-associated molecules ICOS and CTLA4 (**Fig. 3K**).

**Figure 3.**
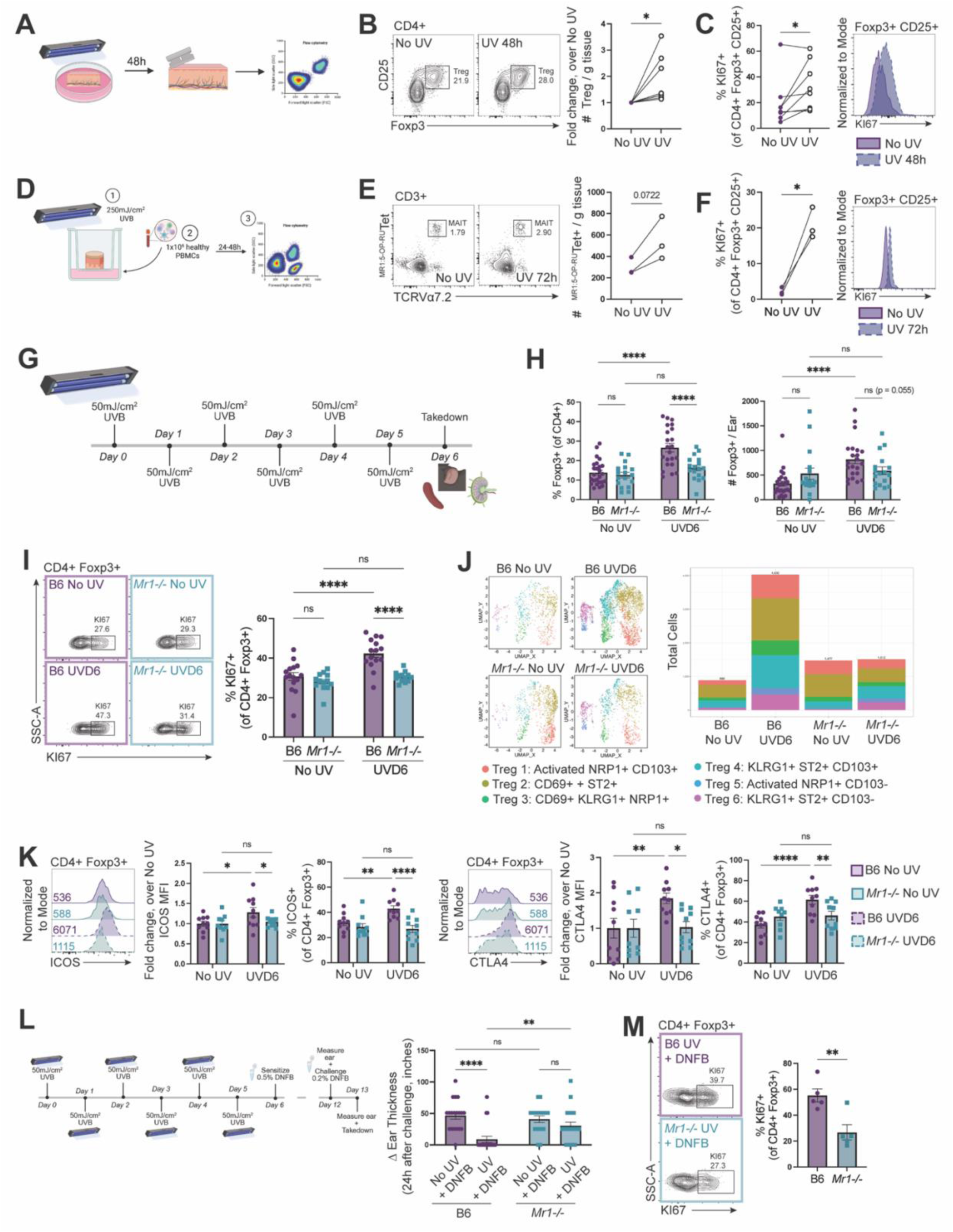
MAIT cells are required for UVB-induced Treg response in healthy skin. (**A**) Schematic of UVB exposure (1×250mJ/cm^2^) of healthy human skin. (**B**) Flow cytometry contour plots of Treg (Live CD45+ CD3+ TCRβ+ CD4+ CD25+ Foxp3+) with and without UVB; Treg number fold change over No UV (n=8/group). (**C**) Flow cytometry histogram of Treg KI67 expression and quantification of KI67+ Treg with and without UVB (n=8/group). (**D**) Schematic of UVB exposure (1×250mJ/cm^2^) of healthy human skin biopsies followed by co-culture with healthy human PBMCs. (**E**) Flow cytometry contour plots and quantification of PBMC-derived MAIT cells (Live CD45+ CD3+ TCRVa7.2+ ^MR1:5-OP-RU^Tetramer+) co-cultured with human skin biopsies with and without UVB (n=3/group). (**F**) Flow cytometry histogram of PBMC Treg KI67 expression and quantification of KI67+ Treg after co-culture with human skin biopsies with and without UVB (n=3/group). (**G**) Schematic of chronic, low-dose UVB in B6 and *Mr1-/-* mice. (**H-I**) Quantification of Treg (H) numbers (live CD45+ γδTCR- CD3+ TCRβ+ CD4+ Foxp3+) and (I) KI67 expression in B6 and *Mr1-/-* skin with (UVD6) and without UVB (No UV) (n=17-24/group). (**J**) Flow cytometry UMAP and corresponding stacked bar plot of Treg from unexposed (No UV) and UVB-exposed (UVD6) B6 and *Mr1-/-* skin (n=5/group). (**K**) Flow cytometry histograms and corresponding quantification of ICOS and CTLA4 on Treg from unexposed (No UV) and UVB-exposed (UVD6) B6 and *Mr1-/-* skin (n=9-10/group). (**L**) Schematic of UVB exposure before induction of delayed contact hypersensitivity (CHS) reactions using the hapten, dinitrofluorobenzene (DNFB); quantification of change in ear thickness over baseline (No UV + vehicle) in B6 and *Mr1-/-* mice after DNFB challenge with (UV) or without (No UV) UVB (n=18-20/group). (**M**) Flow cytometry contour plots and quantification of KI67+ Treg in B6 and *Mr1-/-* skin after UV and DNFB challenge (n=5/group). Data represented as means +/- SEM. Data analyzed with (B-F) paired *t-*test, (H, I, K-L) two-way ANOVA, or (M) Welch’s *t-*test. *p<0.05, **p<0.01, ***p<0.001, ****p<0.0001. ns, not significant.

Given the altered phenotype of Treg in MAIT cell-deficient skin after UVB, we hypothesized that Treg in *Mr1*-/- skin may lack immunosuppressive function. To test this, we compared UVB-mediated suppression of contact hypersensitivity (CHS) in B6 and *Mr1*-/- skin using the established DNFB sensitization-challenge model, in which UVB-induced immunosuppression is Treg dependent (*46–50*). Mice received chronic low-dose UVB on the back skin before DNFB sensitization and a subsequent distal ear challenge; CHS severity was quantified as an increase in ear thickness measured in a blinded fashion (**Fig. 3L**). DNFB sensitization and challenge elicited CHS, i.e. increase in ear thickness, in both B6 and *Mr1*-/- mice (**Fig. 3L**). Consistent with prior studies (*46–50*), UVB markedly reduced CHS in B6 skin, as evidenced by a diminished increase in ear thickness in the UV + DNFB group compared to the DNFB-only group (**Fig. 3L**). However, UV-mediated suppression was lost in *Mr1-/-* mice, which developed ear swelling despite UVB exposure (**Fig. 3L**). Notably, Treg proliferation was reduced in the skin of UV + DNFB-treated *Mr1*-/- mice, compared to MAIT-sufficient B6 controls (**Fig. 3M**).

### MAIT cells suppress photosensitivity in lupus-like skin and promote UV-induced Treg activation in human skin

In contrast to its immunosuppressive effects in healthy skin, UV is immunostimulatory in inflammatory skin diseases such as lupus, in which it can trigger or exacerbate disease (*51–55*). Although impaired Treg responses have been implicated in UV-induced lupus flares in one murine model of disease (*56*), the mechanisms underlying this dysfunctional response to barrier injury remain unclear. Therefore, we next asked how MAIT cell and Treg responses to UVB differ in lupus-like versus healthy skin. Using the photosensitive MRL/lpr model (*57*), we quantified these populations before and after chronic low-dose UVB exposure (as in Fig. 3G). Whereas UVB stimulated MAIT cell expansion in B6 skin, MAIT cell frequency and number were significantly lower in MRL/lpr skin after UVB (**Fig. 4A-B**, fig. S7A-B). Treg expansion was similarly impaired in MRL/lpr skin (**Fig. 4C**, fig. S7A, C). Strikingly, in the absence of UVB-driven MAIT cell and Treg expansion, CD8+ T cell numbers increased ∼30-fold in MRL/lpr but not B6 skin (**Fig. 4D**, fig. S7A). Consistent with these findings, bulk RNA-seq from a small human phototesting study showed that MAIT cell and Treg gene signature scores increased 24 hours after UVB in healthy, but not lupus, skin (fig. S7D-E, **Table 2**).

**Figure 4.**
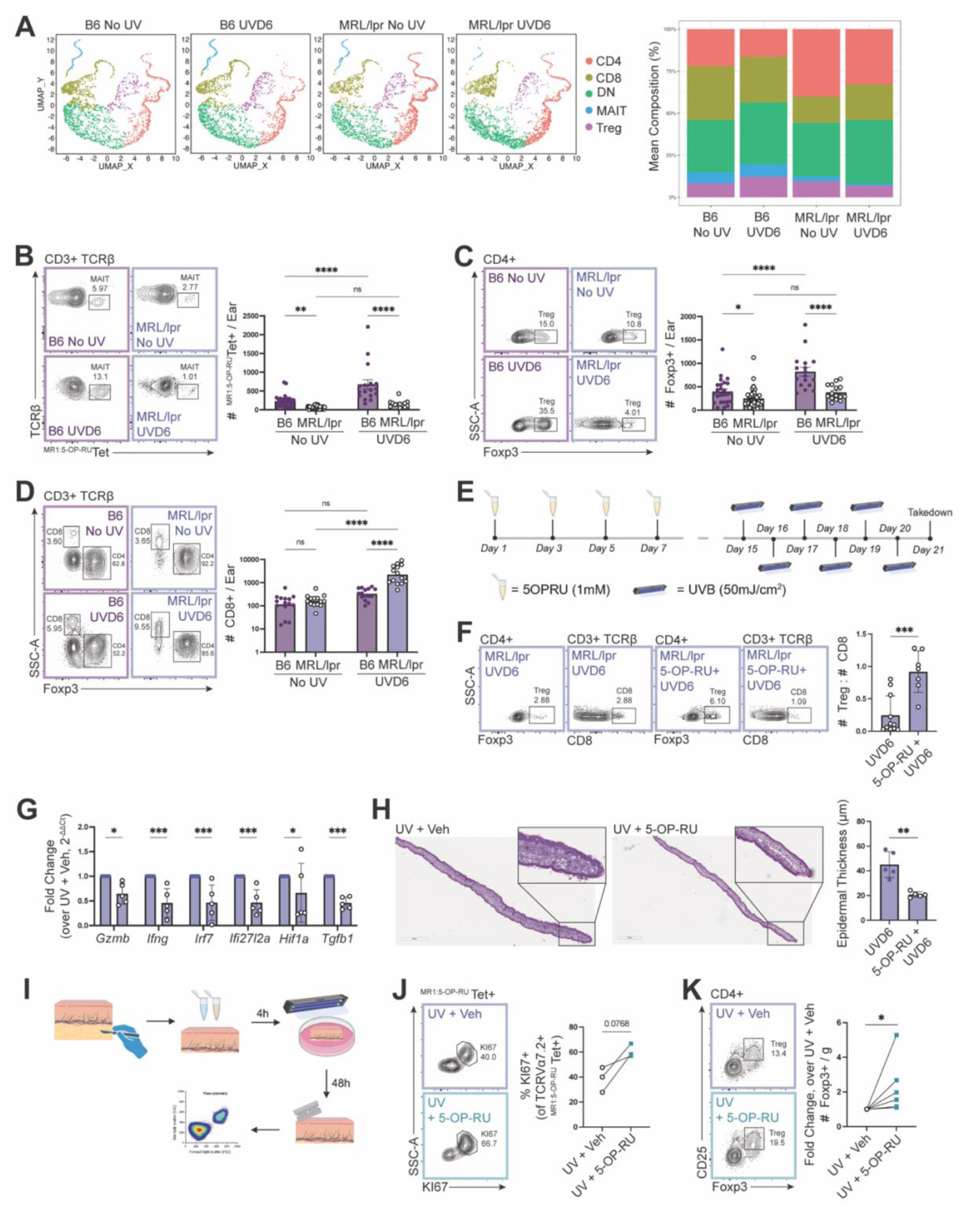
The UVB-induced MAIT and Treg response is impaired in MRL/lpr skin but can be restored by topical 5-OP-RU, and enhanced in human skin. (A) Flow cytometry UMAPs of T cells and corresponding stacked bar plot from unexposed (No UV) and UVB-exposed (UVD6) B6 and MRL/lpr skin (n=5/group). (**B-D**) Flow cytometry contour plots and corresponding quantification of (B) MAIT cells (live CD45+ γδTCR- CD3+ TCRβ+ ^MR1:5-OP-RU^Tetramer+), (C) Treg (live CD45+ γδTCR- CD3+ TCRβ+ CD4+ Foxp3+), and (D) CD8+ T cells in unexposed (No UV) and UVB-exposed (UVD6) B6 and MRL/lpr skin (n=14-16/group). (**E**) Schematic of topical MAIT cell expansion prior to UVB in MRL/lpr mice (female, 8wks). (**F**) Flow cytometry contour plots of Treg and CD8+ T cells, and quantification of Treg:CD8 ratio in UVB-exposed MRL/lpr skin with (UV + 5-OP-RU) or without (UV + Veh) MAIT cell expansion (n=7-11/group). (**G**) Expression of cytotoxic (*Ifng, Gzmb*), interferon-stimulated (*Irf7, Ifi27l2a*), and disease-associated (*Hif1a, Tgfb1*) genes determined by qPCR relative to *Gapdh* in UVB-exposed MRL/lpr skin with (UV + 5-OP-RU) and without (UV + Veh) MAIT cell expansion (n=5/group). (**H**) Representative H&E and quantification of epidermal thickness of UVB-exposed MRL/lpr skin with (UV + 5-OP-RU) and without (UV + Veh) MAIT cell expansion. (**I**) Schematic of MAIT cell expansion with topical 5-OP-RU prior to UVB (1x, 250mJ/cm^2^) in human skin explants. (**J**) Flow cytometry contour plots and quantification of MAIT cell (Live CD45+ CD3+ TCRβ+ TCRVa7.2+ ^MR1:5-OP-RU^Tetramer+) KI67 expression in UVB-exposed human skin with (UV + 5-OP-RU) and without (UV + Veh) 5-OP-RU treatment (n=3/group). (**K**) Flow cytometry contour plots and quantification of Treg (Live CD45+ CD3+ TCRβ+ CD4+ Foxp3+) in UVB-exposed human skin with (UV + 5-OP-RU) and without (UV + Veh) MAIT expansion (n=6/group). Data represented as means +/- SEM. Data analyzed with (B-D) two-way ANOVA, (F-H) Welch’s *t-*test, or (J-K) paired *t-*test. *p<0.05, **p<0.01, ***p<0.001, ****p<0.0001. ns, not significant.

Given the impaired MAIT cell and Treg response to UVB in lupus-like skin, we next tested whether topical antigen-mediated MAIT cell expansion before UVB exposure could attenuate the inflammatory response to this disease-relevant stimulus (**Fig. 4E**). Whereas UVB failed to elicit Treg expansion and instead promoted CD8+ T cell accumulation in vehicle-treated MRL/lpr skin (**Fig. 4C-D**), topical 5-OP-RU treatment significantly increased the Treg:CD8 ratio after UVB (**Fig. 4F**). Consistent with reduced inflammatory activation, 5-OP-RU–treated skin showed lower expression of cytotoxic (*Ifng, Gzmb*), interferon-stimulated (*Irf7, Ifi27l2a*), and disease-associated (*Hif1a, Tgfb1*) transcripts than vehicle-treated skin after UVB (**Fig. 4G**). These immunologic changes were accompanied by reduced epidermal thickness with topical 5-OP-RU (**Fig. 4H**), supporting an immunoregulatory effect of MAIT cell expansion in UVB-induced skin inflammation.

Having established in mice that MAIT cells support cutaneous Treg expansion in response to UVB-driven barrier injury, we next asked whether activating MAIT cells similarly amplifies the UVB-induced Treg response in human skin. Human skin explants were pretreated with 5-OP-RU or vehicle, exposed to acute UVB, and analyzed by flow cytometry 48 hours later (**Fig. 4I**). In the three donors with sufficient MAIT cell recovery, we showed that 5-OP-RU stimulated MAIT cell proliferation (**Fig. 4J**). Across all donors, 5-OP-RU enhanced UVB-induced Treg expansion beyond that seen with UVB alone (**Fig. 4J-K**), indicating that MAIT cells also potentiate UVB-associated Treg responses in human skin. These findings identify a conserved MAIT-Treg axis in cutaneous injury responses that can be engaged to restore the immunoregulatory effects of UVB and prevent injury in inflamed skin.

### CCR2+ monocyte-derived APCs and IL-15 signaling link MAIT cell activation to Treg expansion

Given that MAIT cell-dependent effects on Treg have not been described, we first aimed to identify soluble mediators that could link MAIT cell activation to Treg expansion. Systemic administration of 5-OP-RU induced MAIT cell expansion across tissues, including the skin, and Treg expanded in parallel, recapitulating the effects of topical 5-OP-RU (**Fig. 2B-D**, fig. S8A-G). Luminex profiling identified IL-15 as the only Treg-relevant cytokine significantly increased in the serum of 5-OP-RU-treated mice (**Fig. 5A**). IL-15 is also induced in human skin after UVB (*58*), a context in which we observed MAIT cell-dependent Treg expansion (**Fig. 3A-C**). Therefore, we hypothesized that IL-15 mediates the Treg response downstream of MAIT cells. To test this, we treated B6 and *Il15-/-* mice with topical 5-OP-RU or vehicle (as in Fig. 2A) or exposed them to chronic low-dose UVB (as in Fig. 3G) and quantified skin MAIT cells and Tregs by flow cytometry. Although MAIT cell expansion was preserved in *Il15-/-* skin under both stimuli, Treg accumulation was lost, indicating IL-15 is required for Treg expansion downstream of MAIT cell activation (**Fig. 5B-C**, fig. S9A-B). This defect in Treg response was likely not due to loss of an antigen-presenting myeloid cell; cDC1s were reduced in *Il15-/-* skin only after 5-OP-RU, but not UVB (fig. S10A).

**Figure 5.**
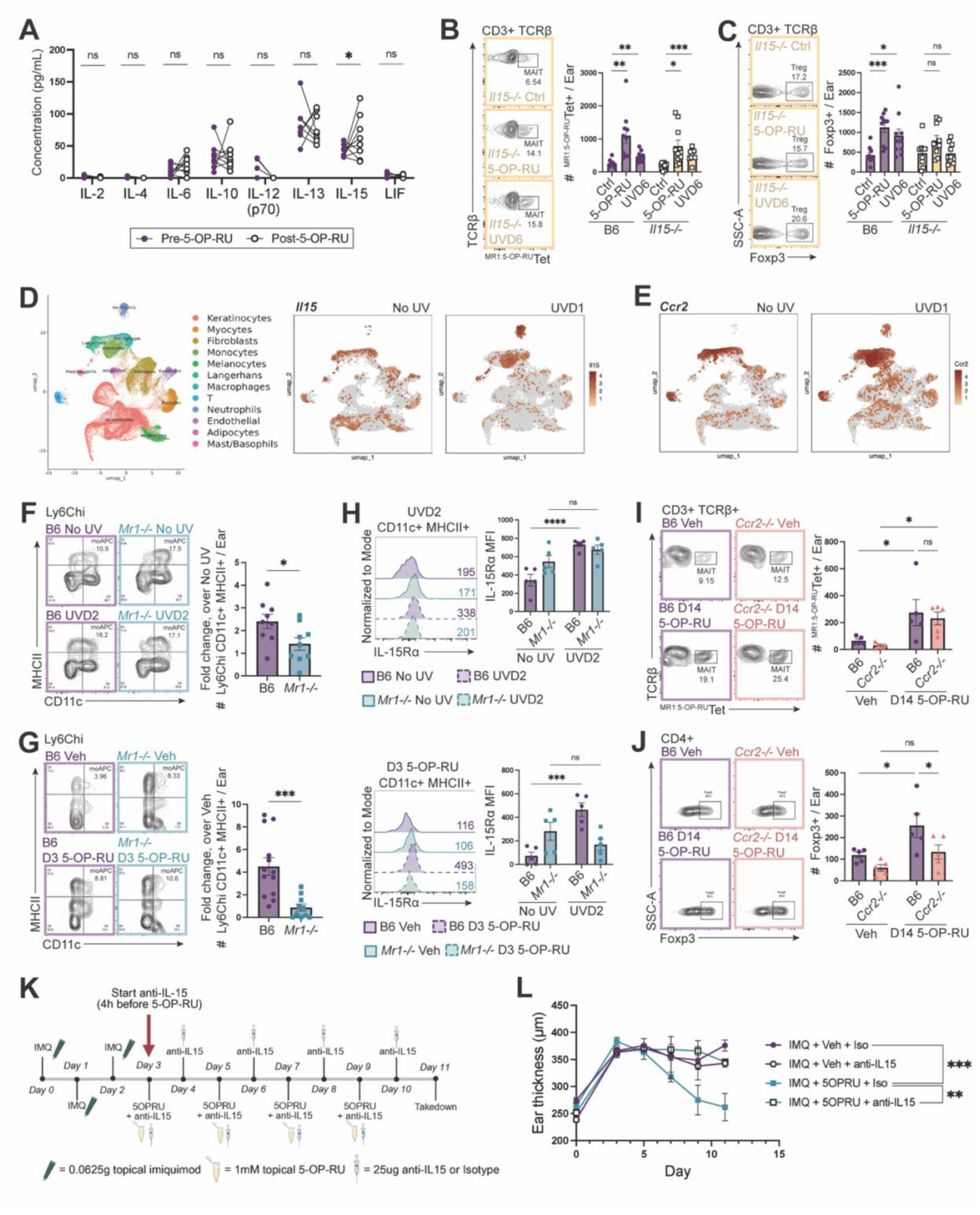
CCR2+ monocyte-derived APCs and IL-15 mediate MAIT cell-driven Treg expansion in the skin. (**A**) Serum cytokine levels before (pre) and after (post) systemic 5-OP-RU (n=8/group). (**B-C**) Flow cytometry contour plots and corresponding quantification of (B) MAIT cells (live CD45+ γδTCR- CD3+ TCRβ+ ^MR1:5-OP-RU^Tetramer+) and (C) Treg (live CD45+ γδTCR-CD3+ TCRβ+ CD4+ Foxp3+) in B6 and *Il15-/-* skin treated with UVB (UVD6) or MAIT cell antigen (5-OP-RU) compared to untreated or vehicle-treated (Ctrl) controls (n=10/group). (**D-E**) ScRNAseq UMAP of stromal and immune cells from untreated (No UV) or UVB-exposed (UV D1) B6 skin and corresponding feature plots of *Il15* and (E) *Ccr2* expression. (**F-G**) Flow cytometry contour plots and corresponding quantification of monocyte-derived antigen-presenting cells (Live CD45+ F4/80- Ly6Chi CD11c+ MHCII+; moAPC) one day after two (F) UVB (UVD2) or (G) 5-OP-RU (D3 5-OP-RU) doses in B6 and *Mr1-/-* skin, relative to untreated or vehicle-treated skin (n=10-13/group). (**H**) Flow cytometry histograms and corresponding quantification of skin moAPC IL15Ra expression after two UVB (UVD2) or 5-OP-RU (D3 5-OP-RU) doses (n=5/group). (**I-J**) Flow cytometry contour plots and corresponding quantification of (I) MAIT cells and (J) Treg in B6 and *Ccr2-/-* skin treated with 5-OP-RU or vehicle (n=5/group). (**K**) Schematic of acute inflammation via topical imiquimod (IMQ) prior to topical 5-OP-RU or vehicle +/- IL-15 blocking IgG in B6 mice. (**L**) Ear thickness measurements over the course of inflammation with IMQ and treatment with 5-OP-RU or vehicle +/- IL-15 IgG (n=5/group). Data represented as means +/-SEM. Data analyzed with (A, F-G) Welch’s *t-*test, (B-C, H-J) two-way ANOVA, or (L) one-way ANOVA. *p<0.05, **p<0.01, ***p<0.001, ****p<0.0001. ns, not significant.

Because MAIT cells are not known to produce IL-15, we next sought to identify the cellular source of IL-15 that links MAIT cell activation to Treg expansion. Single-cell RNA-seq analysis of skin one day after UVB showed that *Il15* is expressed primarily in myeloid cell clusters (**Fig. 5D**), rather than in keratinocytes or fibroblasts, which have been reported to express IL-15 (*59, 60*). *Ccr2*-expressing monocyte-like APCs (*Itgam, Csf1r, Cd14, Itgax, H2-Aa, H2-Ab1*) and neutrophils, both recruited after UVB, expressed high levels of *Il15* (**Fig. 5D-E**, fig. S11). To identify the relevant myeloid population, we asked whether either of these subsets expanded preferentially in B6 compared to *Mr1*-/- skin early after UVB or 5-OP-RU, i.e., before Treg accumulation but during MAIT cell expansion and Treg proliferation (fig. S12). After two doses of either 5-OP-RU or UVB, monocyte-derived APCs (Ly6C^hi^CD11c^+^MHCII^+^; moAPCs) were the only myeloid population to expand in a MAIT-dependent manner with both stimuli, i.e., expanded in B6 but not *Mr1*-/- skin (**Fig. 5F-G**, fig. S13). Neither LCs, conventional DCs, nor neutrophils expanded with 5-OP-RU (fig. S13F, G, I). Although several myeloid subsets upregulated IL-15Rα, neutrophils expressed little surface IL-15Rα despite high *Il15* transcript levels, whereas moAPCs showed high *Il15* transcript levels and marked upregulation of IL-15Rα after UVB or 5-OP-RU (**Fig. 5H**, fig. S14). Importantly, IL-15Rα induction on moAPCs was absent in *Mr1*-/- skin (**Fig. 5H**). Depleting neutrophils did not prevent UV-induced MAIT or Treg expansion, further supporting that neutrophils are not the source of IL-15 required for MAIT-dependent Treg response (fig. S15).

Having identified moAPCs as the likely source of IL-15 downstream of MAIT cells, we next tested whether they are required for MAIT cell-driven Treg expansion in skin. Because MAIT cell and Treg expansion after UVB or 5-OP-RU persisted despite treatment with S1P receptor antagonist FTY720 (fig. S16), indicating local expansion independent of lymph node trafficking, we reasoned that blocking moAPC recruitment to the skin would be sufficient to assess their role. As moAPCs, but not neutrophils, expressed high levels of CCR2 (fig. S17A), we stimulated MAIT cells with 5-OP-RU in *Ccr2-/-* mice, where recruitment of moAPCs, but not other myeloid subsets, is impaired (fig. S17B-F). CCR2 deficiency did not affect 5-OP-RU-induced MAIT cell activation but abrogated Treg expansion (**Fig. 5I-J**). These data support a model in which MAIT cell activation promotes expansion or recruitment of moAPCs, which drive IL-15-dependent Treg proliferation.

Finally, to directly test if the immunoregulatory effects of MAIT cells in inflamed skin require IL-15 signaling, we induced ear inflammation in B6 mice with imiquimod and blocked IL-15 during 5-OP-RU treatment using a neutralizing antibody (**Fig. 5K**). IL-15 blockade abrogated the protective effects of MAIT cells, leaving ears thick and inflamed, whereas inflammation resolved in 5-OP-RU- and isotype-treated controls (**Fig. 5L**). These findings demonstrate that activation of the MAIT-IL-15-Treg axis drives the resolution of established lupus-like skin inflammation.

## Discussion

As in other inflammatory diseases, MAIT cells are reduced in the circulation of patients with chronic inflammatory skin conditions, including systemic sclerosis, dermatomyositis, and lupus (*22–25*). However, due to difficulty obtaining large skin biopsies, low skin MAIT cell abundance, and the technical challenges associated with recovering viable MAIT cells from human skin tissue, very few studies of human skin MAIT cells have been conducted, and the transcriptomic analyses of skin tissues were unsuccessful in reliably capturing MAIT cells (*29–31*). Using cutaneous lupus as a model of chronic skin inflammation sensitive to UV-induced barrier injury, we demonstrate that MAIT cells act as key regulators of cutaneous immune suppression and define a functional MAIT-Treg axis mediated by moAPCs and IL-15. Stimulation with the MR1 ligand 5-OP-RU drove accumulation of both MAIT cells and Treg in healthy and non-lesional lupus-like skin, resolved severe lesions, and suppressed UV-induced inflammation. In the absence of MAIT cells, neither cognate antigen nor UV light induced cutaneous Treg accumulation, activation, or suppressive function. These findings reveal a previously unrecognized immunoregulatory MAIT cell axis in the skin and redefine our understanding of how these cells may contribute to autoimmune and inflammatory skin diseases.

Furthermore, our studies suggest that MAIT cell function is heterogeneous and shaped by the mode of activation and tissue context. Recent *in vitro* studies support this notion, showing that TCR-dependent versus TCR-independent (cytokine-driven) activation induces distinct transcriptional programs and effector functions in MAIT cells (*35, 36*). However, these studies disagree on which mode of activation engages the tissue repair program. A third study by du Halgouet et al. identified basal expression of a tissue repair program in MAIT cells *in vivo* that promoted wound closure independently of TCR engagement (*20*). Together, these studies support a tissue-repair function of MAIT cells but are largely limited to *in vitro* systems or wound models. Our work expands on these findings by showing *in vivo* that TCR-dependent MAIT cell activation can also engage reparative functions, both at homeostasis and in inflamed skin tissue, where MAIT cells are exposed to diverse cytokine cues. The tissue-protective role of MAIT cells has recently been recognized in inflammatory bowel disease and multiple sclerosis (*61, 62*), although the immunomodulatory effects of MAIT cells were not explored.

A central insight from our study is that MAIT cells promote cutaneous Treg responses, revealing an unexpected cellular interaction that supports local immune balance. At homeostasis, skin Treg maintain tolerance to commensal microbes, promote stem cell cycling, and orchestrate wound repair (*63–65*). Decades of work have shown that UVB induces local and systemic Treg activation in healthy skin, leading to immune suppression (*46–50, 66–68*). Here, we show that UVB elicits coordinated expansion of MAIT cells and Treg in healthy skin and that genetic loss of MAIT cells abrogates the Treg response and impairs UV-induced immunosuppression. UV is also a potent trigger of lupus disease (*33*). Although alterations in Treg number and function have been described in lupus (*69–72*), only one prior study implicated defective UV-elicited Treg response in lupus-prone mice, attributing this to elevated IFN-I in the skin (*56*). In stark contrast to healthy skin, our findings reveal that both MAIT cells and Treg fail to expand in response to UVB in lupus-prone skin. Instead, CD8+ T cells accumulate, consistent with inflammatory and cytotoxic responses to UVB-induced hypoxia (*73, 74*). Thus, impaired MAIT cell responsiveness may represent an upstream contributor to Treg dysfunction and photosensitivity in lupus. Notably, our studies showed that targeted MAIT cell activation restores the UVB-induced Treg response and suppresses inflammation in lupus-like skin, highlighting the therapeutic potential of engaging the MAIT-Treg axis to re-establish immune balance in inflamed tissue.

Mechanistically, MAIT cell activation stimulated production of IL-15. Though IL-15 is often regarded as a growth and survival signal for CD8+ T cells, several studies have demonstrated its capacity to promote Treg differentiation and proliferation (*59, 75–78*), supporting its potential role in the cutaneous MAIT-Treg axis. IL-15 signaling characteristically occurs in *trans,* whereby membrane-bound IL-15Rα on the surface of one cell presents IL-15 to the shared IL-2/IL-15 receptor on the receiving cell (*79*). Although keratinocytes, fibroblasts, and several dendritic cell subsets (Langerhans cells (LCs) and monocyte-derived dendritic cells (moDCs)) can produce IL-15 (*59, 60, 77*), MAIT cell activation selectively upregulated IL-15Rα on the surface of moAPCs and, to a lesser extent, neutrophils, nominating moAPCs as the likely candidate for IL-15 trans-presentation to Treg. Consistent with this, CCR2 deficiency prevented moAPC recruitment into the skin following MAIT cell activation and abrogated downstream Treg expansion. Together, these data support a model in which MAIT cells license moAPCs to trans-present IL-15 to Treg, thereby promoting tissue-protective immune regulation in skin.

Our study focused on tissue-protective mechanisms downstream of MAIT cells in the skin, particularly those driving Treg responses, but several questions remain. First, the novel repair cascade elucidated in the present study begins at the MAIT cells, but how MAIT cells become activated remains to be explored. Recent work by Haas et al. showed cell-type-specific MR1 upregulation in APCs and stromal cells depending on the nature of the microbial infection (*80*), suggesting that topical 5-OP-RU may be presented by different MR1-expressing cells in the context of a healthy versus an inflamed skin environment. The upstream signals that activate MAIT cells after UV also remain unknown. UV-induced stress may promote MAIT cell activation through non-mutually exclusive mechanisms, including altered microbial metabolites, endogenous MR1 ligands, or cytokine-driven activation independent of classical antigen presentation. IFN-I is a candidate trigger, as UV induces IFN-I production (*56, 81–83*), and MAIT cells can be activated via IFN-I receptor (*9, 10*). UV also perturbs the skin microbiome and its metabolic milieu (*84, 85*), potentially altering the availability of MAIT cell antigens. Defining these pathways will clarify how environmental stress engages MAIT cells in the skin.

Finally, although MAIT cell activation drives downstream Treg expansion and function, whether this Treg response requires antigen-specific stimulation or instead depends on cytokine-or contact-mediated signals remains unresolved. Prior studies implicate LCs and CD80/86 expression in UVB-induced Treg response (*47, 86*), and antigen-specific Treg expansion has been observed after UV in the presence of an exogenous antigen (*50, 87*). It also remains unclear how MAIT cells support moAPCs upstream of Treg expansion. MAIT cells can promote differentiation of monocyte-derived dendritic cells through CD40-CD40L interactions (*88*), and MAIT cell production of GM-CSF, IFN-γ, and TNF-α can support APC function of other myeloid cells (*89*). Defining the antigen dependence of this pathway and the mechanisms by which MAIT cells license moAPCs will be important for understanding how immune regulation and tissue repair are coordinated in the skin.

Overall, this work identifies MAIT cells as key orchestrators of skin immune homeostasis and repair, acting through moAPC and IL-15 signaling to drive protective Treg responses. These findings broaden current understanding of MAIT cell biology at barrier tissues and suggest that strategies to enhance MAIT cell function or restore MAIT–Treg crosstalk may help promote tissue repair and limit immunopathology in autoimmune skin disease.

## Materials and Methods

### Mice and *in vivo* lupus models

MRL/lpr and *Ccr2-/-* mice were purchased from Jackson Laboratories (Strains 00485 and 004999, respectively). *Mr1-/-* mice were obtained from Dr. Michael Constantinides, and wildtype littermates were used as controls (B6). *Il15-/-* mice were obtained from Dr. Mary Jo Turk. All mice were at least 8 weeks old at the start of the experiment, and controls were age- and sex-matched. All mice were maintained in a specific pathogen-free facility at the Geisel School of Medicine at Dartmouth. All studies were approved by Dartmouth College’s Institutional Animal Care and Use Committee. To induce lupus-like inflammation in the ears, 0.06-0.07g imiquimod cream (5%) was applied to the ears daily for three total applications. Inflammation was confirmed by measuring ear thickness.

### MAIT cell activation via 5-OP-RU

5-OP-RU for topical and systemic activation of MAIT cells was generated by combining 1mM 5-Amino-6-(D-ribitylamino)uracil (hydrochloride) (5-A-RU-HCl, Cayman) with 2-methylglyoxal (Sigma) in a 50-fold molar excess. For topical activation, the mixture was applied to the ears (∼75 ul per ear) of mice with a cotton-tipped applicator every 48 hours for a total of 4 (healthy B6 skin, non-lesional MRL/lpr skin, and IMQ-treated skin) or 9 (spontaneous lesion treatment in MRL/lpr mice) doses. For systemic activation, the mixture (200 ul) was injected intraperitoneally every day for 3 total doses, then a 3-day rest period, followed by an additional round of 3 daily injections. For human studies, a single dose of 5-OP-RU was applied topically to cover large skin tissue explants for 4 hours prior to UV exposure.

### Histology

Murine ear or back skin was fixed in 10% Neutral Buffered Formalin for 24 hours, then paraffin-embedded. Formalin-fixed, paraffin-embedded tissue sections (4μm) were air-dried at room temperature before loading on the Tissue-Tek Prisma Stainer (Sakura Finetek USA) in the Dartmouth Health Pathology Shared Resource. Automated H&E protocol includes bake and dewax, hematoxylin, clarifier, bluing agent, and eosin followed by dehydration in graded alcohols, clearing in xylene, and automated coverslip application. Reagents used are as follows: Hematoxylin 2 (Richard-Allan Scientific), Clarifier 2 (Richard-Allan Scientific), Bluing Agent (Richard-Allan Scientific), Eosin-Y (Richard Allan Scientific).

### RNA Isolation and qPCR

RNA from whole skin was isolated using the RNeasy Fibrous Tissue kit (Qiagen). Reverse transcription was performed using the iScript cDNA synthesis kit (Bio-rad) and quantitative PCR executed with iTaq Universal SYBR Green Supermix (Bio-rad). Primer sequences are in Table 3. Gene expression CT values were normalized to *Gapdh*.

### Flow Cytometry

Ear pinnae were excised and separated into dorsal and ventral sheets. Sheets were digested dermal side down in RPMI 1640 containing Liberase TL (0.25mg/mL, Roche) and DNase I (0.5mg/mL, Sigma) at 37°C for 1 hour and 45 minutes. Ears were then minced with microscissors and transferred to gentleMACS C tubes (Miltenyi) for mechanical dissociation. Dissociated tissue was passed through a 70 μm filter and washed with PBS. Single cell suspensions were stained with Live Dead blue viability dye (ThermoFisher), ^MR1:5-OP-RU^Tetramer (NIH Tetramer core), TruStain FcX PLUS (BioLegend, 1:100), and antibody cocktail. Cells were fixed for intracellular staining using Foxp3/Transcription Factor Fix/Perm (Cytek). For cytokine production assays, single cell suspensions were incubated with Brefeldin A (BioLegend) and Cell Stimulation Cocktail (ThermoFisher) for 6 hours at 37°C before proceeding with staining. Flow cytometry was performed using a Cytek Aurora 657. FlowJo software (Treestar Incorporated) was used to analyze data. Flow cytometry UMAPs were generated using SpectroFlo.

### Single cell RNA sequencing

Single-cell RNA sequencing was performed using the Chromium GEM-X Single Cell 5’ Reagent Kit v3 (10x Genomics) according to the manufacturer’s instructions (CG000684). Briefly, cells were loaded onto the 10x Genomics Chromium Controller to generate single-cell GEMs, followed by barcoding, cell lysis, and cDNA synthesis. Gene expression and V(D)J libraries were prepared in parallel from the same GEM reaction. Following cDNA amplification, gene expression libraries were constructed using the Chromium Single Cell 5’ Library Kit (CG000684), and T cell receptor (TCR) V(D)J libraries were prepared using the Chromium Single Cell Mouse TCR Amplification Kit according to the manufacturer’s protocol (CG000698). All libraries were quality-assessed by Agilent TapeStation and prior to sequencing. Gene expression and V(D)J libraries were sequenced on an Illumina NextSeq 2000 instrument targeting a minimum of 30,000 reads per cell for gene expression libraries and 5,000 reads per cell for V(D)J libraries, using recommended read configurations (28 bp Read 1, 10 bp i7 index, 10 bp i5 index, 90 bp Read 2).

### Human skin processing for immune cells

Healthy human surgical skin discards were obtained from the Dartmouth-Hitchcock Medical Center Pathology (IRB exempt). Subcutaneous adipose tissue was removed by scraping. Skin tissue was arranged dermal-side down and cultured in 1:1 DMEM/F-12 (Fisher) supplemented with 10% fetal bovine serum (Fisher) at room temperature, with the epidermal surface maintained above the air-liquid interface. To isolate immune cells, the upper layer of skin was removed with a razor blade, minced into fine pieces, then digested in RPMI 1640 (Fisher) supplemented with Collagenase IV (1mg/mL; Sigma) and DNase I (2mg/mL; Sigma) for 1 hour at 37°C with shaking. Cell suspensions were then passed through a 70µm filter, washed with PBS, and counted.

### UVB irradiation

For UVB irradiation, mice were anesthetized with isoflurane and exposed to 50mJ/cm^2^ UVB every 24 hours for 6 total doses using Tyler Research UV-2 irradiation and dosimetry system. Human skin tissue was exposed to a single 250mJ/cm^2^ dose of UVB and processed 48 hours later.

### Human skin explant and PBMC co-cultures

Healthy surgical discards were prepared as above. Punch biopsies (6mm) were placed epidermal side up in a Transwell insert, then exposed to a single acute dose of UVB (250mJ/cm^2^). Immediately following UVB exposure, 1 million peripheral blood mononuclear cells (PBMCs) isolated from healthy donors (DH IRB, STUDY02001542) were added to the well beneath the skin explant in 350 ul. PBMCs were maintained in T Cell Media (RPMI 1640 supplemented with 10% fetal bovine serum, 1% Penicillin/Streptomycin, 1mM L-glutamine, 1mM sodium pyruvate, 1X MEM non-essential amino acids, 1X HEPES, 50μM β-mercaptoethanol). Co-cultures were incubated at 37°C for 72 hours, then PBMCs were collected and stained for flow cytometry as above.

### Contact hypersensitivity assays

Five days prior to UVB exposure, an approximately 7.5cm^2^ area of mouse dorsal skin was shaved with an electric razor. A thin layer of Nair™ was applied for 30 seconds and rinsed off with water. Mice were then exposed (unless no UV controls) to 50mJ/cm^2^ UVB every 24 hours for 6 total doses using Tyler Research UV-2 irradiation and dosimetry system. 24 hours after the final UVB exposure, skin was sensitized by swabbing 0.5% dinitrofluorobenzene (DNFB, Sigma) onto the shaved dorsum. Six days later, a baseline measurement of ear thickness was recorded, and the skin was then challenged at a distal site by swabbing 0.2% DNFB or vehicle (4:1 olive oil:acetone) onto the ears. 24 hours after challenge, ear thickness was measured and recorded to assess the presence or absence of a contact hypersensitivity reaction.

### Human phototesting and bulk RNAseq

Human phototesting and subsequent RNA sequencing was completed as previously described in (*90*). MAIT cell and Treg gene signatures were defined as in (*91–94*).

### 10x Flex Genomics Single Cell RNA-sequencing

Formalin fixed paraffin embedded (FFPE) skin tissue blocks were prepared for the 10x Genomics Flex assay according to manufacturer’s protocol (CG000606). 4 x 25μm sections per block were deparaffinized with xylene, rehydrated with progressive ethanol dilutions and dissociated in Liberase TM (Roche) solution on a gentleMACS tissue dissociator (Miltenyi). The resulting suspension was passed through a 70μm strainer and quantified using AO/PI staining on a Luna FX7 automated fluorescent cell counter. The 10x Genomics Flex assay was run per manufacturer’s protocol (CG000527). Briefly, cells were thawed from -80C, counted, and up to 1×10^6^ cells were put into the hybridization reaction using the Mouse Transcriptome Probe Kit containing sample specific barcodes. Pooled samples were loaded onto a 10x Genomics Chromium X instrument, and 10,000 cells per sample were captured. Concentration and size of sequencing libraries were measured by Qubit (ThermoFisher) and Tape Station (Agilent), respectively. Sequencing was performed on a NextSeq2000 instrument (Illumina) targeting 25,000 reads/cell. The cellranger v9.0.0 pipeline was used to process raw fastq files before downstream analysis. Data were analyzed using Seurat (v5.3). Immune and stromal clusters were identified by unsupervised clustering, and cell identity was defined by the top 5 expressed genes in each cluster.

### Statistical Analysis and Data Availability

Statistical comparisons were performed using GraphPad Prism software V.11 (GraphPad Software, La Jolla, CA, USA). The statistical test, representation of statistical significance, and sample sizes are listed in the figure legends for each analysis. Treg 10X scRNAseq data will be available under Gene Expression Omnibus study number GSExxxxxx. 10X Flex scRNAseq data will be publicly available under Gene Expression Omnibus study number GSExxxxxx.

## Ethics Statement

Human samples were obtained under protocols approved by the Dartmouth Health Institutional Review Board (DH IRB, STUDY02001542). All samples were de-identified prior to analysis, and all procedures complied with relevant ethical regulations. All animal experiments were conducted in accordance with institutional guidelines and were approved by the Dartmouth College Institutional Animal Care and Use Committee (IACUC) (protocol number: 2262). All efforts were made to minimize animal suffering and to reduce the number of animals used.

## Author Contributions

S.S.G., G.E.C designed the study. G.E.C., A.A., V.C., Z.T.P., L.O., S.K., L.K.M., O.F.B., K.D., H.B.G., S.L., F.W.K. conducted experiments and/or analyzed data. G.E.C. and S.S.G. wrote the manuscript, and D.B., P.C.R., M.S.S., T.A.L., D.A.R., M.G.C. critically reviewed and edited it.

## Acknowledgements

This work was supported by NIAMS R01AR080641 to S.S.G, T32 AI007363 and the Lupus Foundation of America Gina M. Finzi Memorial Student Summer Fellowship Award to G.C, and Friends of Dartmouth Cancer Center funding (ICI COMPEL award) to S.S.G. and P.C.R. The authors acknowledge the support of the Dartmouth Cancer Center Shared Resources, which are funded in part by the NCI Cancer Center Support Grant P30CA023108. Genomic analyses were performed by the Genomics and Molecular Biology Shared Resource (RRID:SCR_021293), supported by grants S10OD030242, P20GM130454, and S10OD025235. Single-cell and spatial genomics experiments were conducted using the 10x Chromium platform within the same resource. Immune profiling was carried out by the Immune Monitoring and Flow Cytometry Shared Resource (RRID:SCR_019165). Histological analyses were provided by the Pathology Shared Resource (RRID:SCR_023479). The NIH Tetramer Core Facility (NIH Contract 75N93020D00005 and RRID:SCR_026557) provided mouse and human 5-OP-RU-loaded MR1 tetramers. Experiment schematics were created in BioRender.

