## Supplementary Figures and Tables for "Mucosal-associated invariant T cells support IL-15-dependent Treg response in skin injury and promote resolution of skin inflammation"

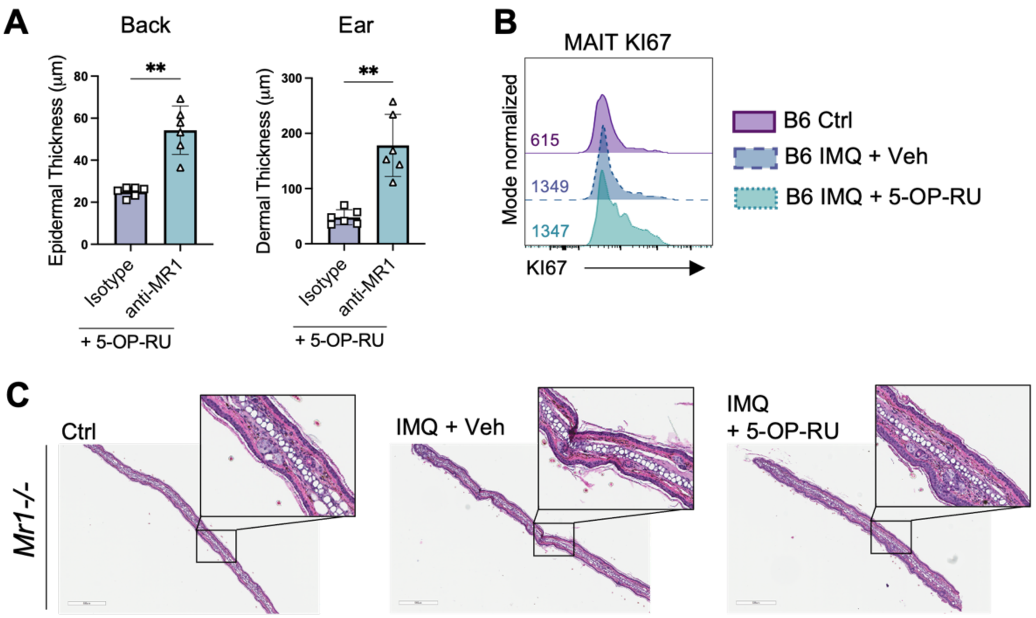


**Figure S1. Lupus-like skin inflammation is resolved by antigen-mediated activation of MAIT cells.** (**A**) Quantification of back epidermal or ear dermal thickness in MRL/lpr mice treated with 5-OP-RU with (anti-MR1) or without (Isotype) MR1 blockade (n=6/group). (**B**) Flow cytometry histogram of MAIT cell KI67 (proliferation) expression in untreated (Ctrl) or Imiquimod (IMQ)-treated B6 skin with (5-OP-RU) or without (Veh) MAIT cell expansion (n=5/group). (**C**) Representative H&E stain of control or imiquimod-treated *Mr1-/-* ears with (5-OP-RU) and without (Veh) MAIT cell expansion. Data represented as mean +/- SEM. Data analyzed with (A) Welch's *t*-test. **p<0.01.

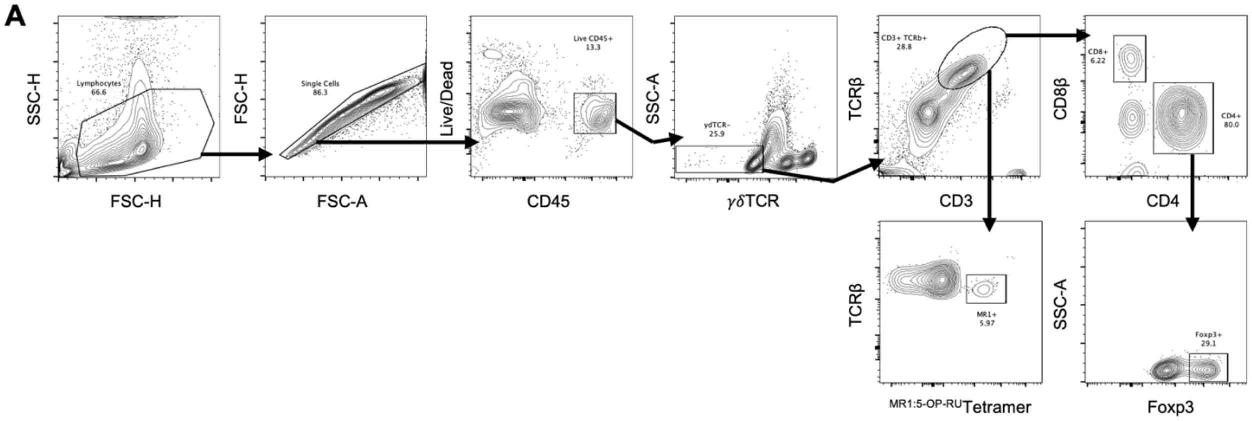


**Figure S2. Gating strategy for skin T cells.** (**A**) Representative flow cytometry gating strategy for identification of T cells in murine skin.

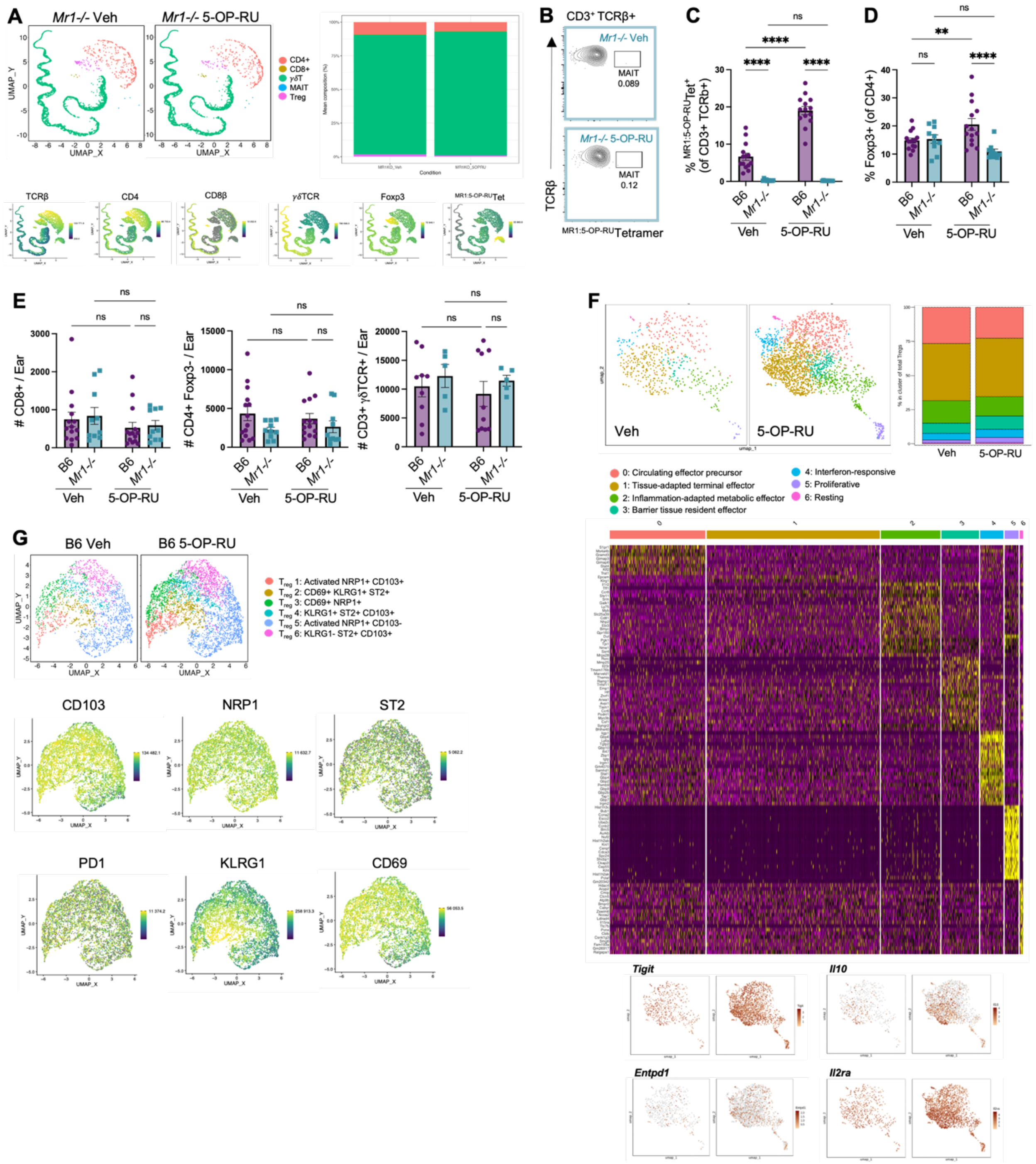


**Figure S3. MAIT cell activation via topical 5-OP-RU drives heterogeneous Treg expansion in healthy skin.** (**A**) Flow cytometry UMAP and corresponding stacked bar plot of T cells with (5-OP-RU) and without (Veh) MAIT cell antigen exposure in *Mr1-/-* skin (n=5/group). (**B**) Flow cytometry contour plots of MAIT cells (live CD45+ γδTCR- CD3+ TCRβ+ $^{\text{MR1:5-OP-RU}}$Tetramer+) in *Mr1-/-* skin with (5-OP-RU) and without (Veh) antigen exposure. (**C-D**) Flow cytometry quantification of (C) MAIT cell and (D) Treg frequency in B6 and *Mr1-/-* skin with (5-OP-RU) and without (Veh) antigen exposure (n=14/group). (**E**) Flow cytometry quantification of skin T cell subsets in B6 and *Mr1-/-* mice with (5-OP-RU) and without (Veh) MAIT cell antigen exposure (n=5-14/group). (**F**) ScRNAseq UMAP and corresponding stacked bar plot, heatmap, and feature plots for Treg captured from skin of 5-OP-RU- or vehicle-treated B6.*Foxp3-GFP* mice. (**G**) Flow cytometry UMAP and corresponding feature plots of Treg from 5-OP-RU- or Vehicle-treated B6 skin (n=5/group). Data represented as means +/- SEM. Data analyzed with (C-E) two-way ANOVA. $^{**}p<0.01$, $^{****}p<0.0001$. ns, not significant.

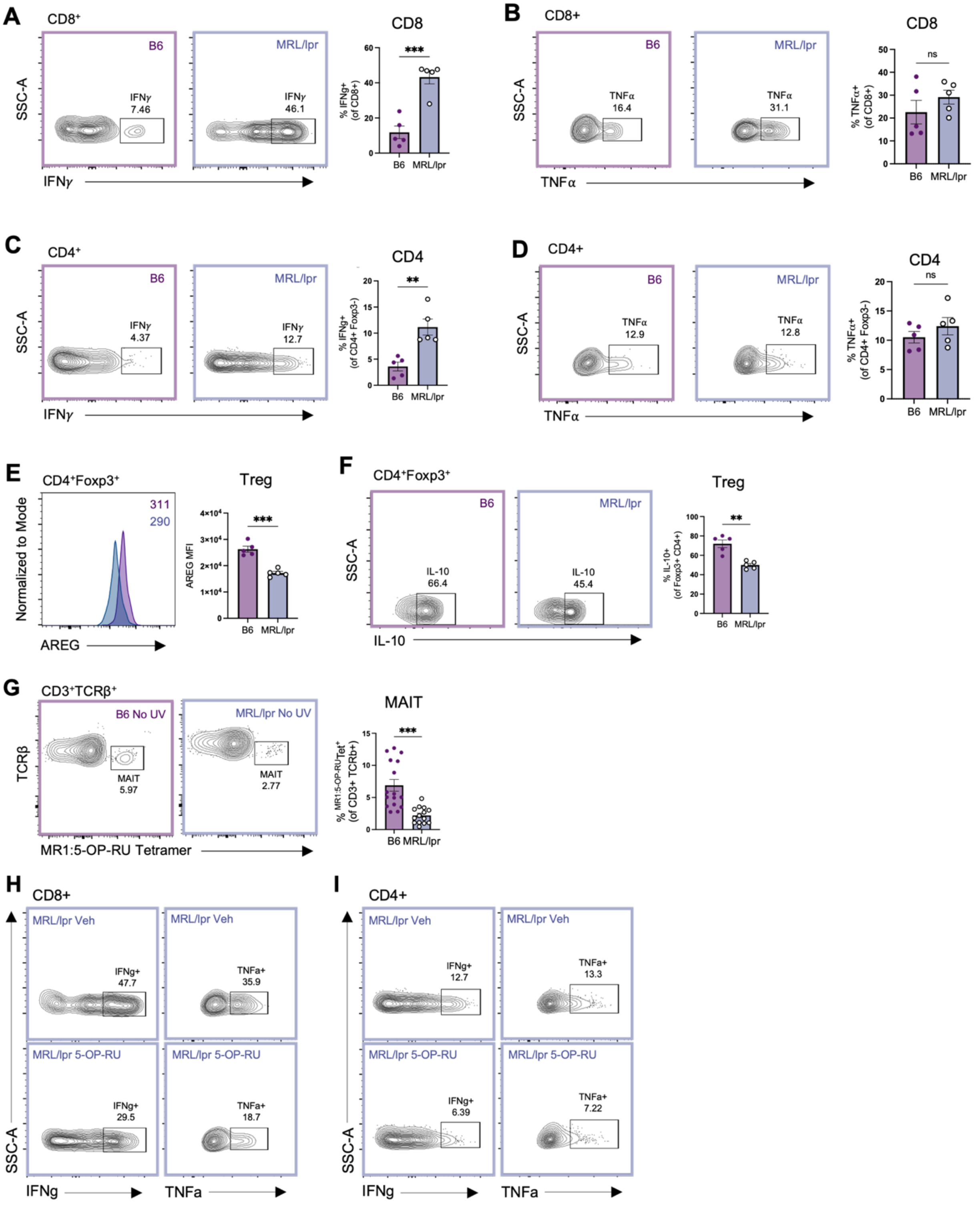


**Figure S4. T cell cytokine production at baseline and following 5-OP-RU treatment in B6 and MRL/lpr non-lesional skin.** (**A-B**) Flow cytometry contour plots and quantification of CD8+ T cell (A) IFN-g and (B) TNF-a production in B6 and MRL/lpr skin at baseline (n=5/group). (**C-D**) Flow cytometry contour plots and quantification of CD4+ T cell (C) IFN-g and (D) TNF-a production in B6 and MRL/lpr skin at baseline (n=5/group). (**E**) Flow cytometry histogram and quantification of Treg AREG expression in B6 and MRL/lpr skin at baseline (n=5/group). (**F**) Flow cytometry contour plot and quantification of Treg IL-10 production in B6 and MRL/lpr skin at baseline (n=5/group). (**G**) Flow cytometry contour plots and corresponding quantification of MAIT cell frequency in B6 and MRL/lpr skin at baseline (n=14-16/group). (**H-I**) Representative flow cytometry gating for IFN-g and TNF-a production by (H) CD8+ and (I) CD4+ T cells in MRL/lpr skin with (5-OP-RU) and without (Veh) MAIT cell stimulation. Data represented as means +/- SEM. Data analyzed with (A-G) *t*-test. **p<0.01, ***p<0.001. ns, not significant.

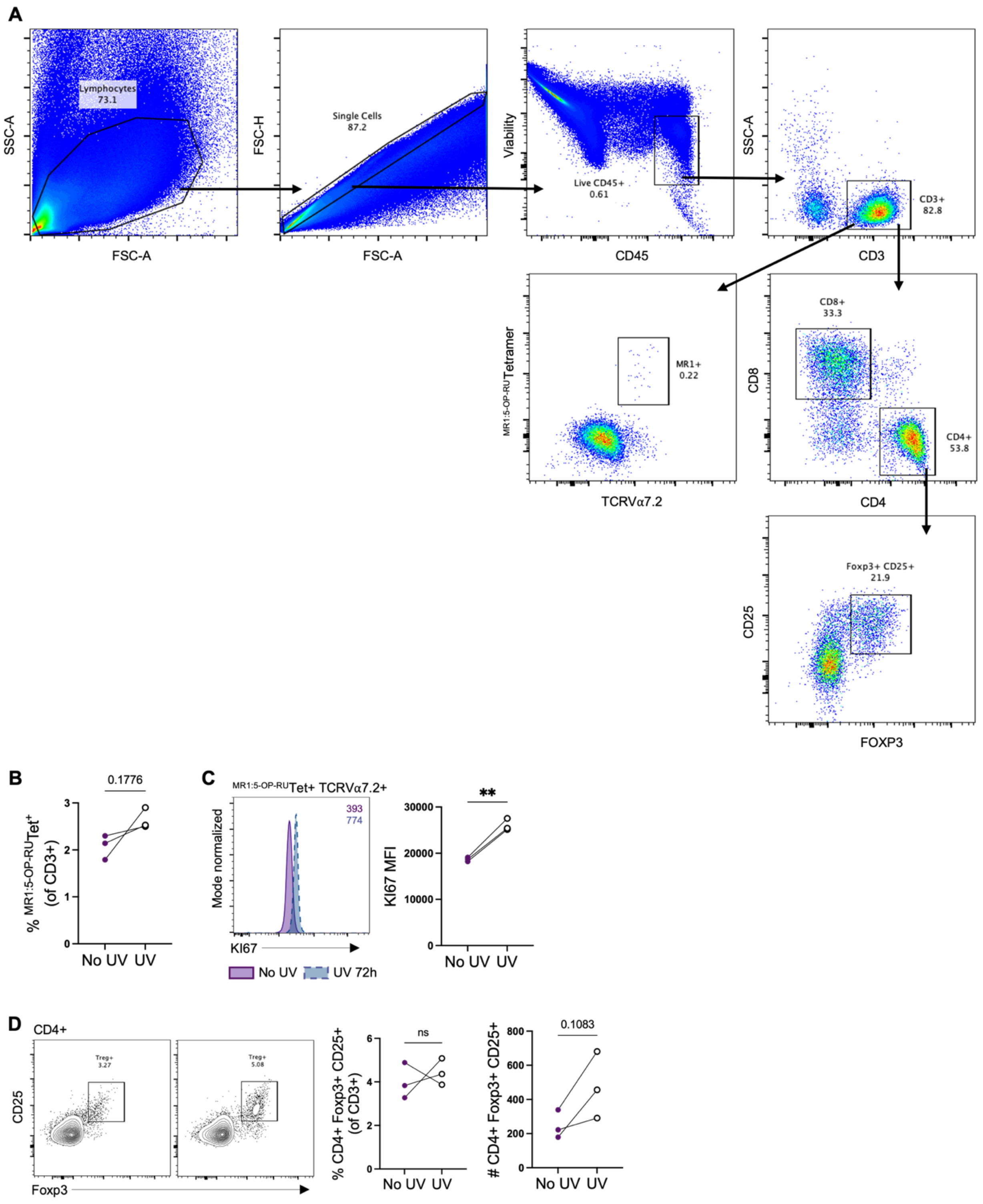


**Figure S5. UVB exposure of human skin drives proliferation and expansion of PBMC-derived MAIT cells and Treg.** (**A**) Representative flow cytometry gating strategy for identification of T cells in human skin. (**B**) Quantification of PBMC-derived human MAIT cells after co-culture with healthy human skin that was either exposed (UV) or not exposed (No UV) to a single, acute dose of UVB (n=3/group). (**C**) Flow cytometry histogram and corresponding quantification of KI67 mean fluorescence intensity on PBMC-derived Treg after co-culture with healthy human skin that was either exposed (UV) or not exposed (No UV) to a single, acute dose of UVB (n=3/group). (**D**) Flow cytometry contour plots and corresponding quantification of PBMC-derived Treg frequency and absolute number after co-culture with healthy human skin that was either exposed (UV) or not exposed (No UV) to a single, acute dose of UVB (n=3/group). Data analyzed with (B-D) paired *t*-test. **$p<0.01$. ns, not significant.

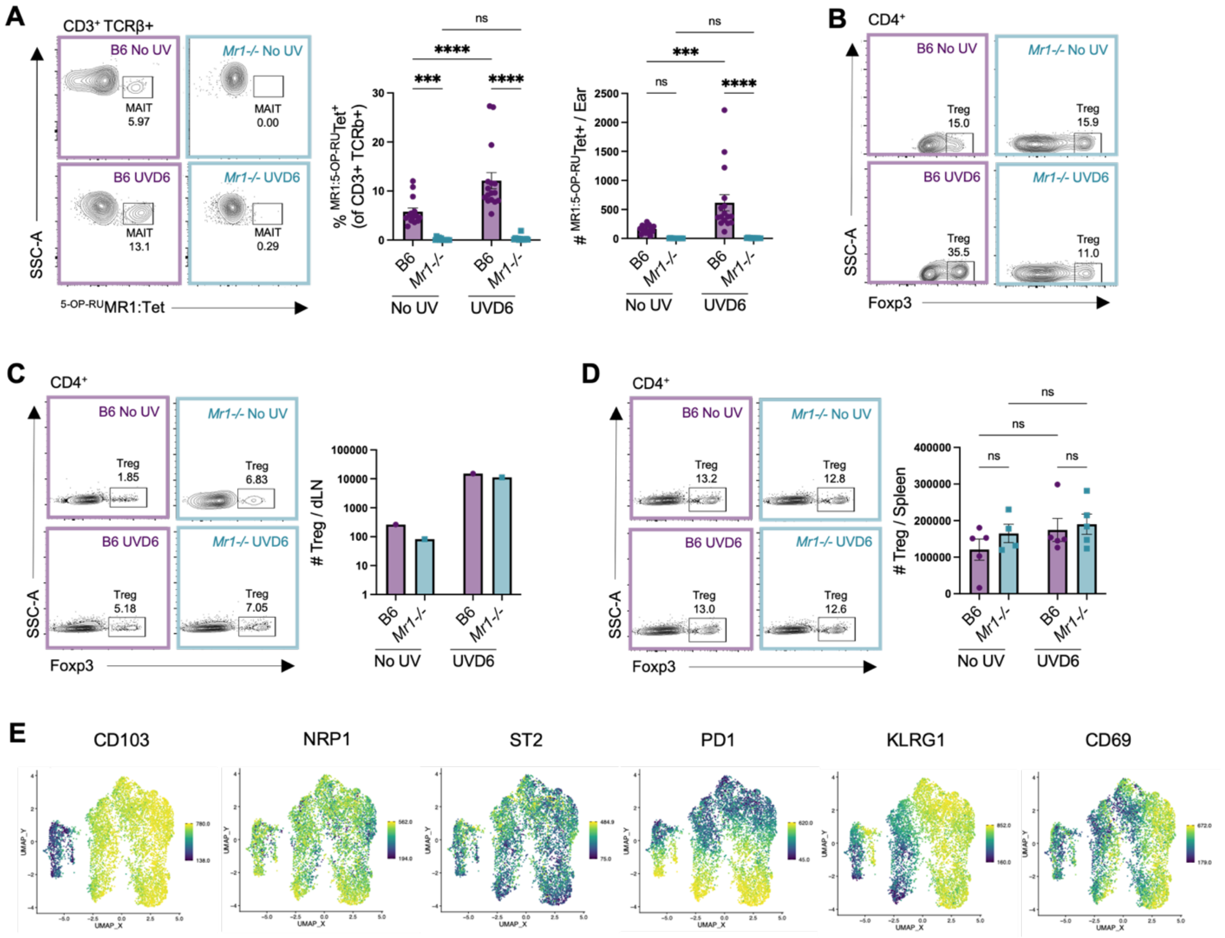


**Figure S6. UVB-induced MAIT cell activation supports heterogeneous skin Treg expansion.** (**A**) Flow cytometry contour plots and corresponding quantification of skin MAIT cells (live CD45+ γδTCR- CD3+ TCRβ+ $^{MR1:5\text{-}OP\text{-}RU}$Tetramer+) in B6 and *Mr1-/-* mice with (UVD6) and without (No UV) chronic UVB exposure (n=17-24/group). (**B**) Flow cytometry contour plots of skin Treg (live CD45+ γδTCR- CD3+ TCRβ+ CD4+ Foxp3+) with (UVD6) and without (No UV) chronic UVB exposure in B6 and *Mr1-/-* skin. (**C**) Flow cytometry contour plots and corresponding quantification of Treg in B6 and *Mr1-/-* skin draining lymph nodes with (UVD6) and without (No UV) UVB exposure. Data concatenated from SDLNs of 5 mice. (**D**) Flow cytometry contour plots and corresponding quantification of Treg in B6 and *Mr1-/-* spleens with (UVD6) and without (No UV) UVB (n=5/group). (**E**) Feature plots of select Treg markers used to subset Treg from flow cytometric data of B6 and *Mr1-/-* mice at baseline (No UV) or after UVB exposure (UVD6). Data represented as means +/- SEM. Data analyzed with (A, D) two-way ANOVA. ***$p<0.001$, ****$p<0.0001$. ns, not significant.

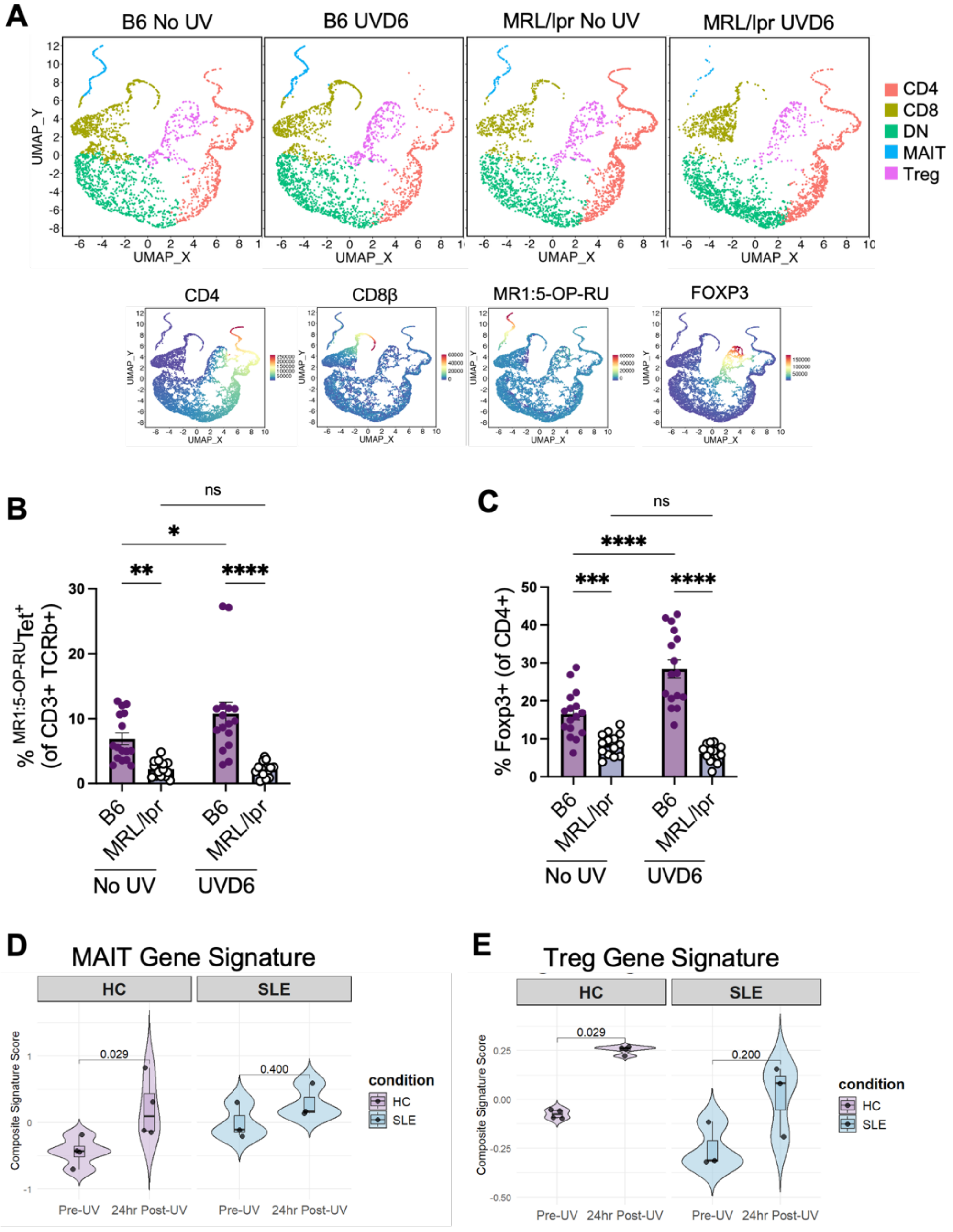


**Figure S7. T cell responses to UVB are altered in lupus skin.** (**A**) Flow cytometry UMAP of skin T cells generated from concatenated flow cytometric data of representative baseline (No UV) and UVB-exposed (UVD6) B6 and MRL/lpr mice (n=5/group). Stacked bar plot quantifying mean composition per T cell cluster. (**B-C**) Quantification of (B) MAIT cell and (C) Treg frequency in B6 and MRL/lpr skin with (UVD6) and without (No UV) UVB. (**D-E**) Expression of (D) MAIT cell- or (E) Treg-associated genes in healthy and lupus skin before (No UV) and 24 hours after acute UVB (2 x MED; n=3-4/group). Data represented as means +/- SEM. Data analyzed with (B, C) two-way ANOVA or (D, E) paired *t*-test. *p<0.05, **p<0.01, ***p<0.001, ****p<0.0001. ns, not significant.

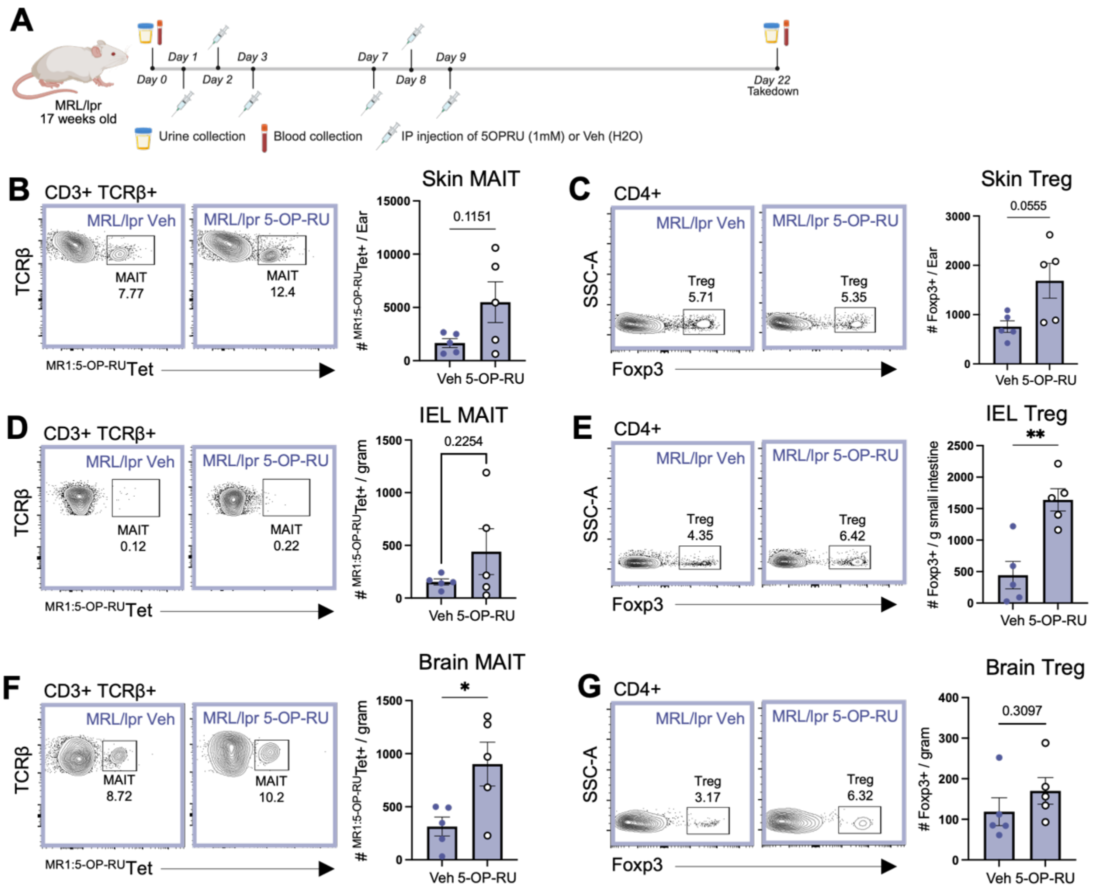


**Figure S8. Systemic 5-OP-RU triggers MAIT cell and Treg expansion across multiple tissues.** (**A**) Schematic of systemic delivery of MAIT antigen 5-OP-RU or vehicle to MRL/lpr mice (16-20wks, female). (**B**-**G**) Flow cytometry contour plots and corresponding quantification of (B, D, F) MAIT cells or (C, E, G) Treg in (B-C) skin, (D-E) intestinal epithelial lymphocytes (IEL), or (F-G) brain with (5-OP-RU) or without (Veh) systemic MAIT cell activation (n=5/group). Data represented as means +/- SEM. Data analyzed with Welch's *t*-test. *p<0.05, **p<0.01. ns, not significant.

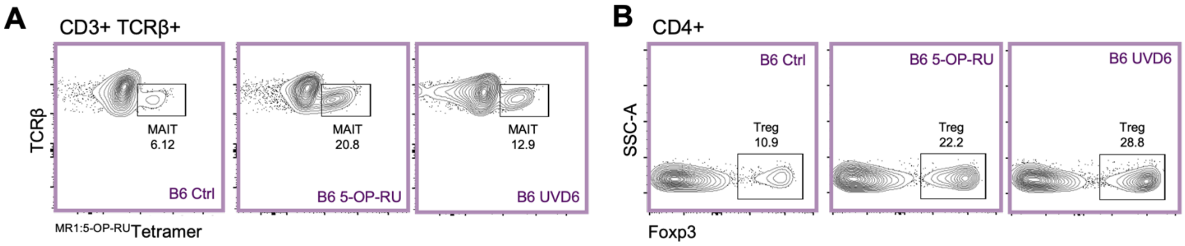


**Figure S9. Gating strategy for B6 MAIT cells and Treg from Figure 5B-C.** (**A-B**) Flow cytometry contour plots of (A) MAIT cells and (B) Treg from B6 skin at baseline (Ctrl) or treated with antigen (5-OP-RU) or UVB (UVD6).

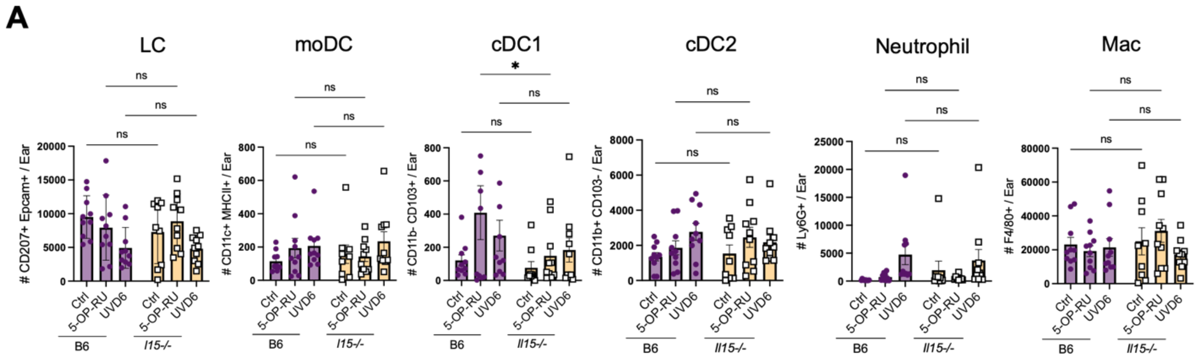


**Figure S10. IL-15 deficiency does not affect myeloid cell numbers in the skin.** (**A**) Quantification of antigen-presenting cells in B6 and *Il15-/-* skin at baseline (Ctrl) or after antigen (5-OP-RU) or UVB (UVD6; n=10/group). Data represented as means +/- SEM. Data analyzed with two-way ANOVA. *p<0.05. ns, not significant. Mac = macrophages

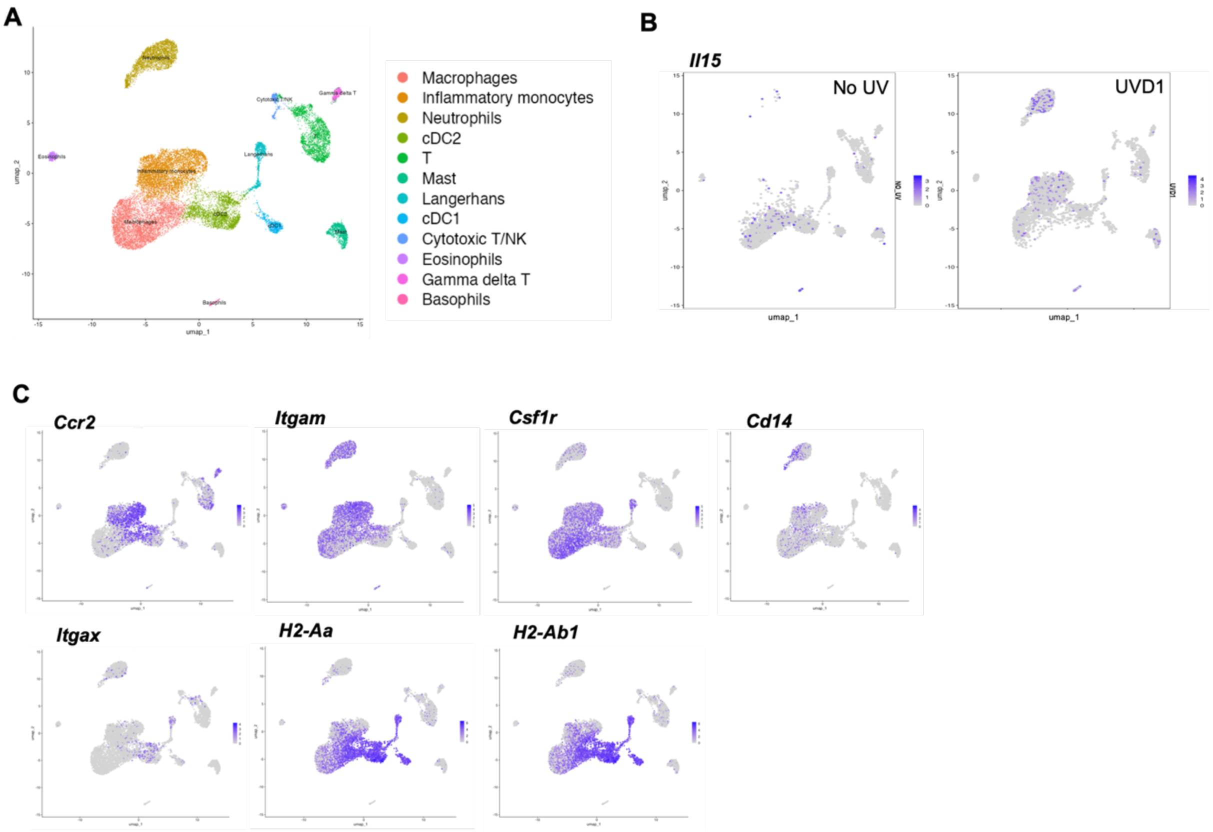


**Figure S11. Monocytes and neutrophils are the primary expressors of *Il15* early after skin UVB exposure.** (**A**) ScRNAseq UMAP of immune cells captured in B6 skin after a single dose of UVB. (**B**) Feature plots of skin immune cell *Il15* gene expression with (UVD1) and without (No UV) UVB. (**C**) Feature plots of skin immune cell moAPC-associated genes (*Ccr2, Cd14, Itgax, H2-Aa, H2-Ab1*) captured by scRNAseq.

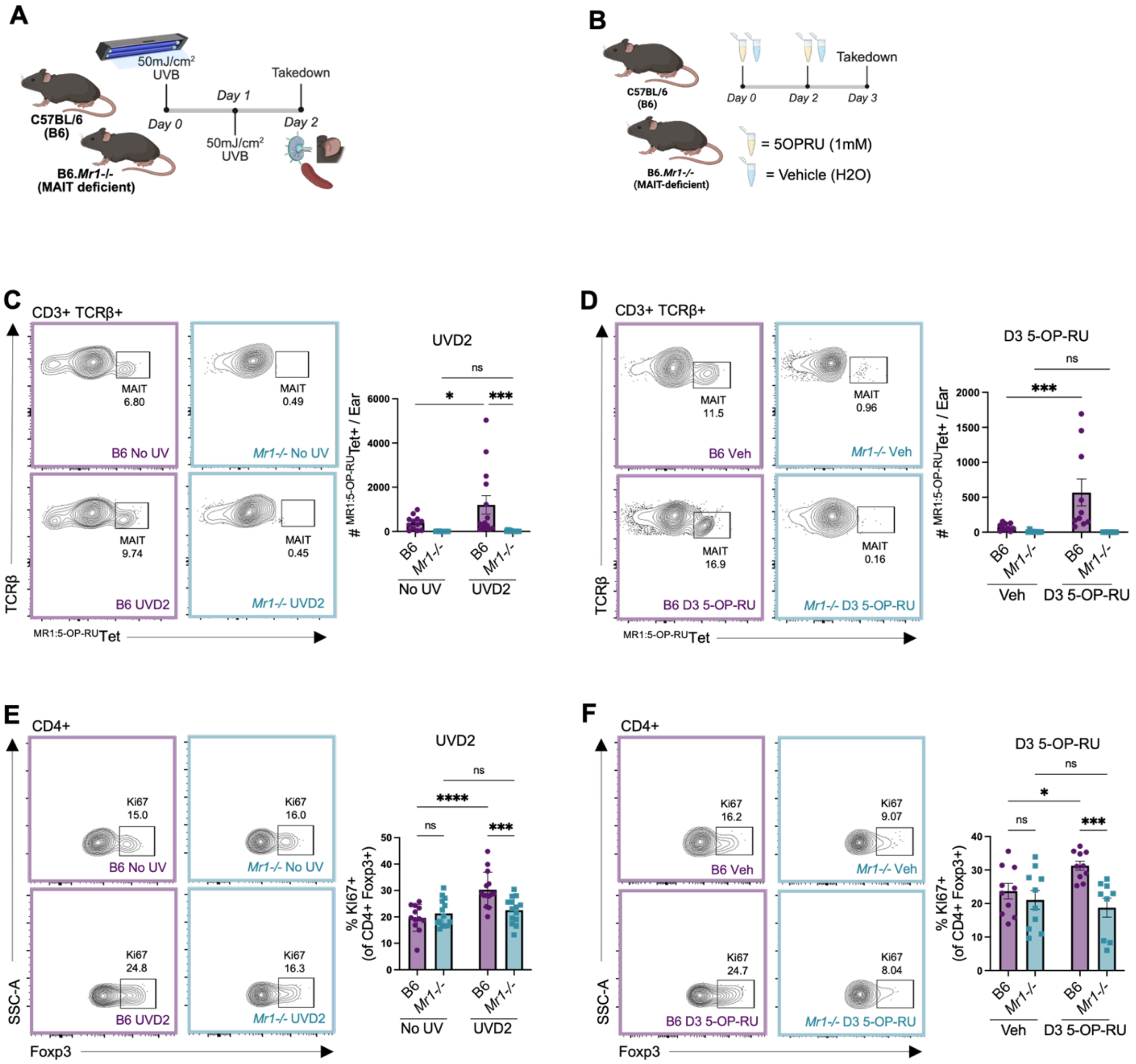


**Figure S12. MAIT cells expand early after UV or 5-OP-RU to support Treg proliferation.** (**A-B**) Schematic of brief MAIT cell stimulation with (A) UVB (2x 50mJ/cm$^2$) or (B) 5-OP-RU (2x 1mM) in B6 and *Mr1-/-* skin. (**C-D**) Flow cytometry contour plots and corresponding quantification of MAIT cells early after (C) UV (UVD2; n=14/group) or (D) 5-OP-RU (D3 5-OP-RU; n=9-10/group). (**E-F**) Flow cytometry contour plots and corresponding quantification of KI67+ Treg early after (C) UV (UVD2; n=14/group) or (D) 5-OP-RU (D3 5-OP-RU; n=9-10/group). Data represented as means +/- SEM. Data analyzed with two-way ANOVA. *p<0.05, ***p<0.001, ****p<0.0001. ns, not significant.

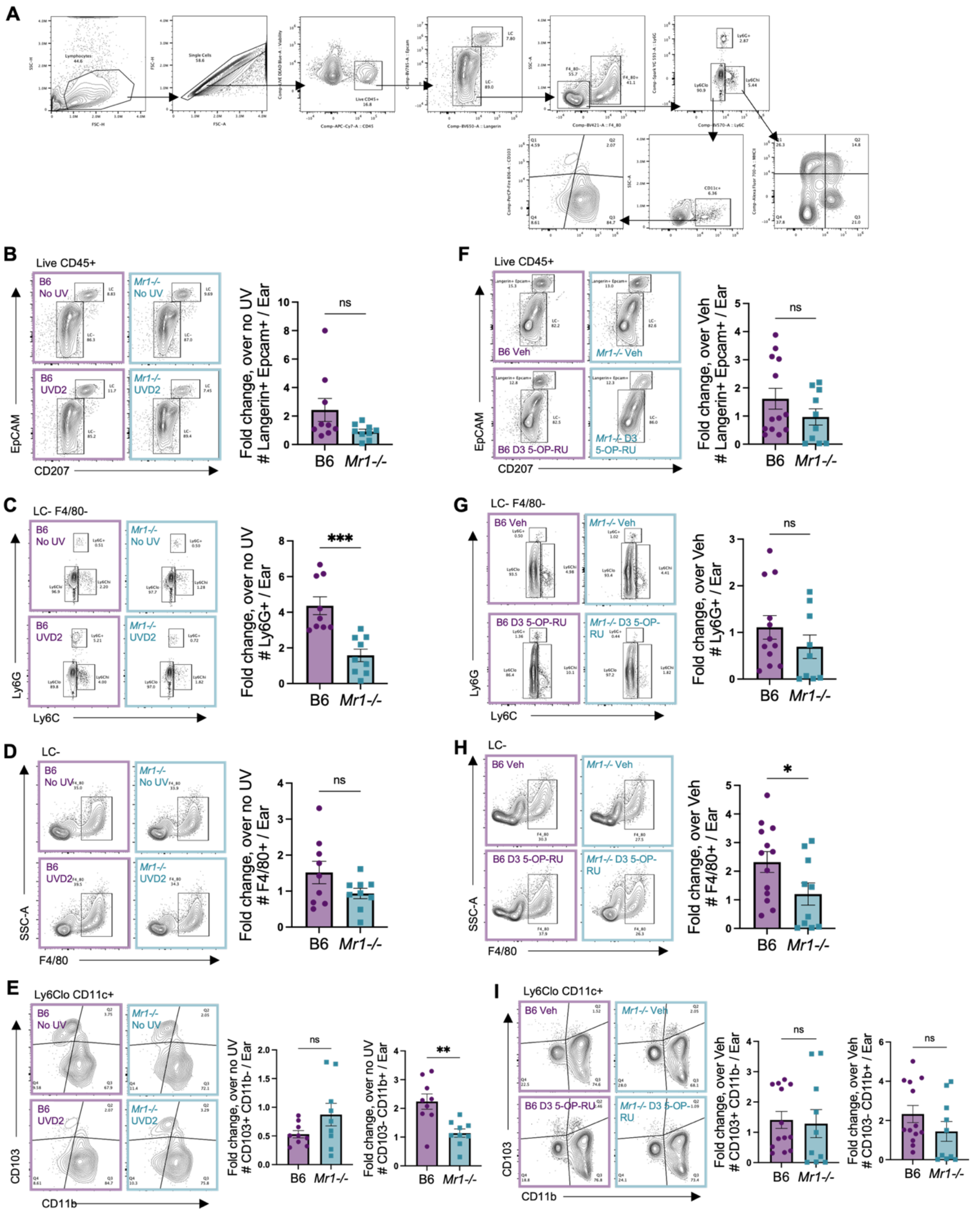


**Figure S13. Myeloid cell expansion early after MAIT cell stimulation.** (**A**) Flow cytometry gating strategy for identification of skin myeloid cell subsets: neutrophils (live CD45+ CD207- EpCAM- F4/80- Ly6G+), cDC1 (live CD45+ CD207- EpCAM- F4/80- Ly6Clo CD11b- CD103+), cDC2 (live CD45+ CD207- EpCAM- F4/80- Ly6Clo CD11b+ CD103-), Langerhans cell (live

CD45+ CD207+ EpCAM+), moAPC (live CD45+ CD207- EpCAM- F4/80- Ly6Chi CD11c+ MHCII+), and macrophages (Live CD45+ CD207- EpCAM- F4/80+). (**B-I**) Representative flow cytometry contour plots and corresponding quantification of (B, F) Langerhans cells, (C, G) neutrophils, (D, H) macrophages, or (E, I) conventional dendritic cells early after (B-E) UV (UVD2) or (F-I) 5-OP-RU (D3 5-OP-RU). Data represented as means +/- SEM. Data analyzed with Welch's *t*-test. *p<0.05, **p<0.01, ***p<0.001. ns, not significant.

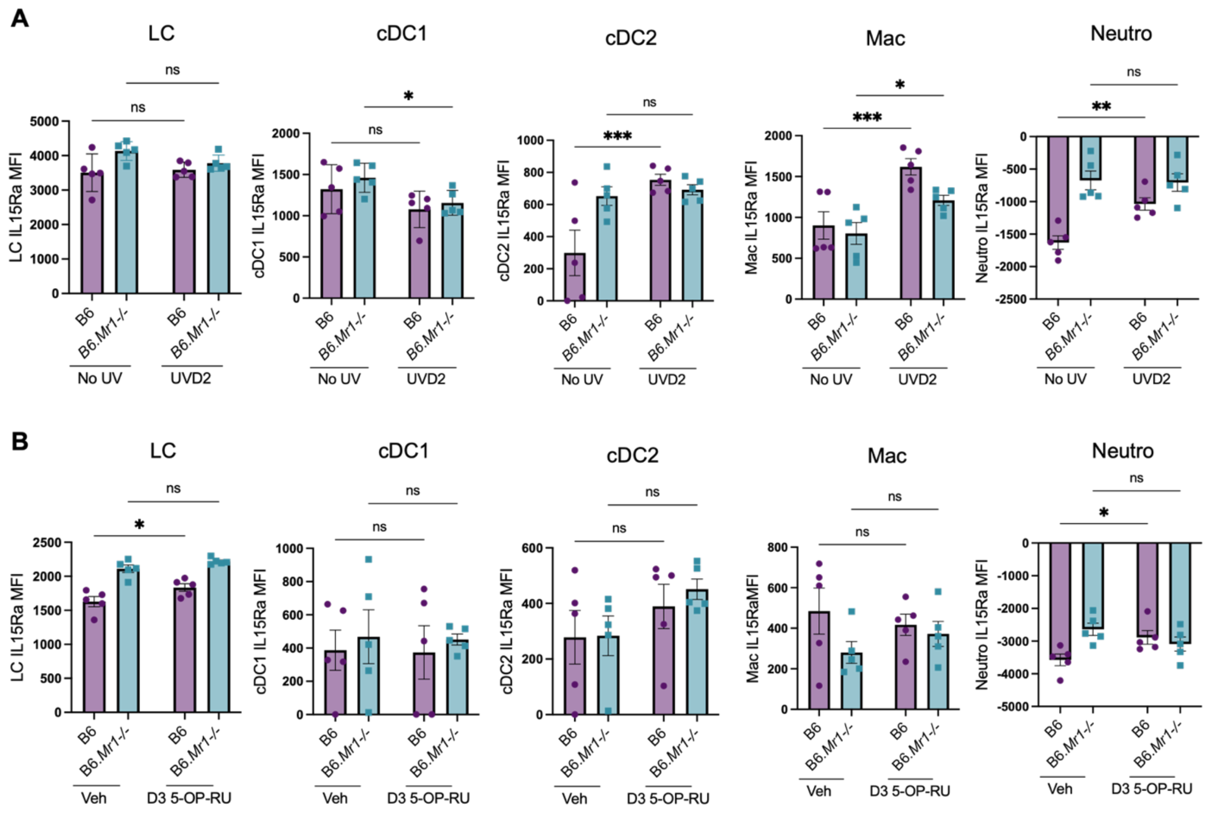


**Figure S14. Induction of IL15Ra expression on myeloid cells after MAIT cell stimulation.** (**A-B**) Quantification of IL15Ra expression on myeloid cell subsets early after (A) UVB (UVD2) or (B) 5-OP-RU (D3 5-OP-RU) in B6 and *Mr1-/-* skin (n=5/group). Data represented as means +/- SEM. Data analyzed with two-way ANOVA. *p<0.05, **p<0.01, ***p<0.001. ns, not significant.

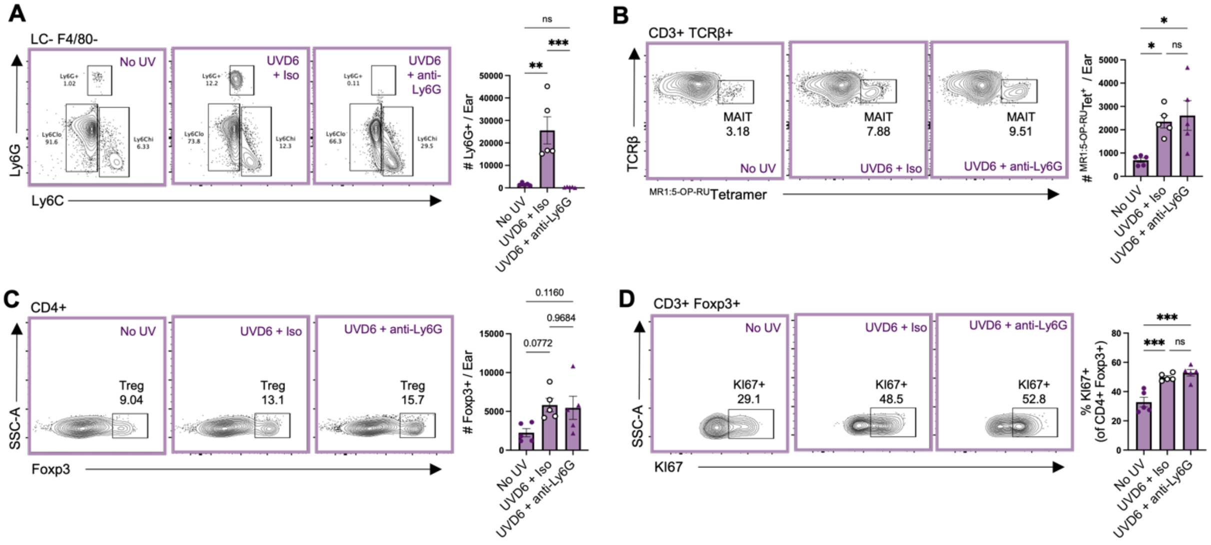


**Figure S15. Neutrophils are not required for UVB-induced Treg expansion and proliferation downstream of MAIT cells.** (**A-D**) Representative flow cytometry contour plots and corresponding quantification of skin (A) neutrophils, (B) MAIT cells, (C) Treg, and (D) frequency of KI67+ Treg at baseline (No UV) or after chronic UVB exposure with (UVD6 + anti-Ly6G) or without (UVD6 + Iso) neutrophil depletion. Data represented as means +/- SEM. Data analyzed with One-way ANOVA. *p<0.05, **p<0.01, ***p<0.001. ns, not significant.

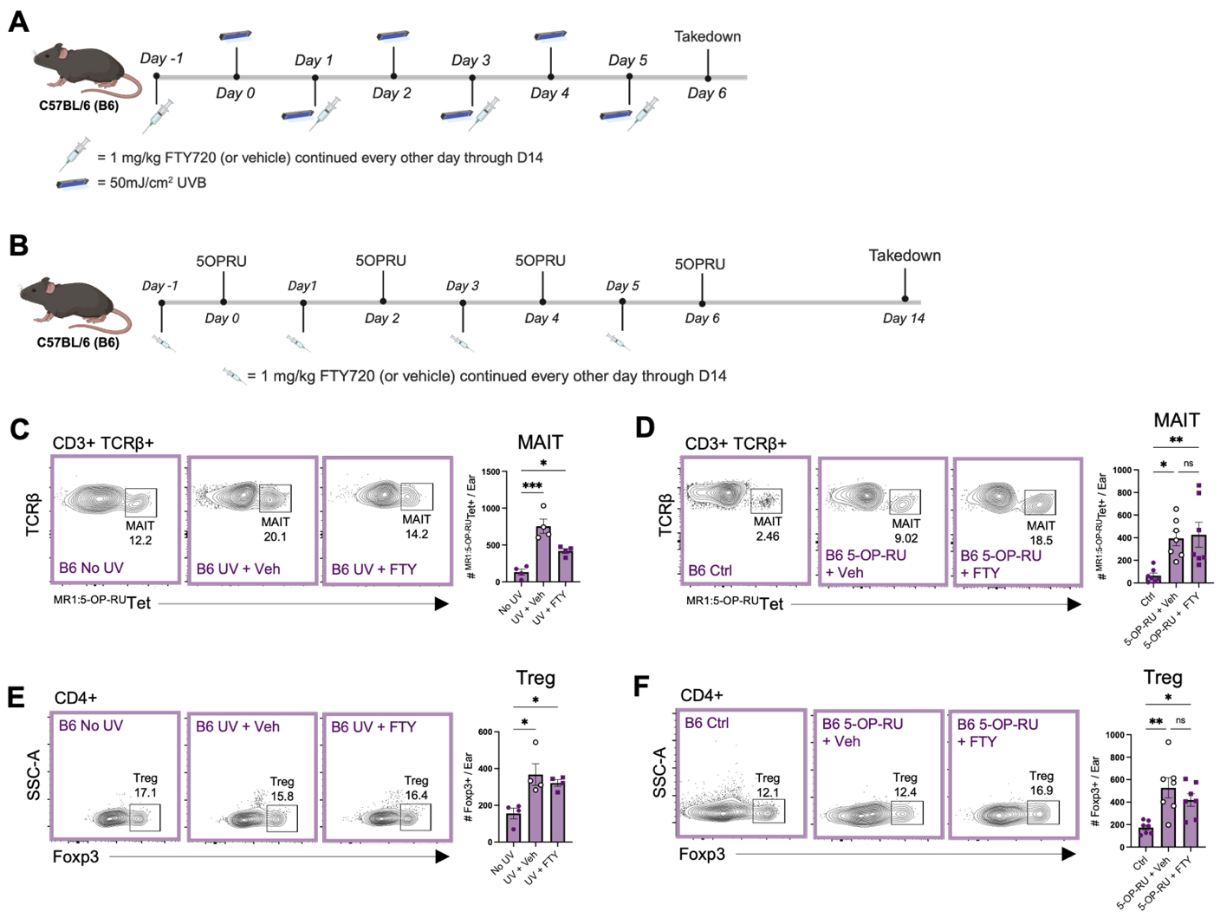


**Figure S16. MAIT cell stimulus drives local expansion of MAIT cells and Treg in the skin.** (**A-B**) Schematic of S1P receptor blockade (FTY720) in B6 mice during exposure to MAIT cell stimuli. (**C-D**) Flow cytometry contour plots and corresponding quantification of skin MAIT cells after (C) UVB (n=4/group) or (D) 5-OP-RU (n=7/group) in the presence or absence of FTY720 (FTY). (**E-F**) Flow cytometry contour plots and corresponding quantification of skin Treg after (E) UVB (n=4/group) or (F) 5-OP-RU (n=7/group) in the presence or absence of FTY. Data represented as means +/- SEM. Data analyzed with one-way ANOVA. *p<0.05, **p<0.01, ***p<0.001. ns, not significant.

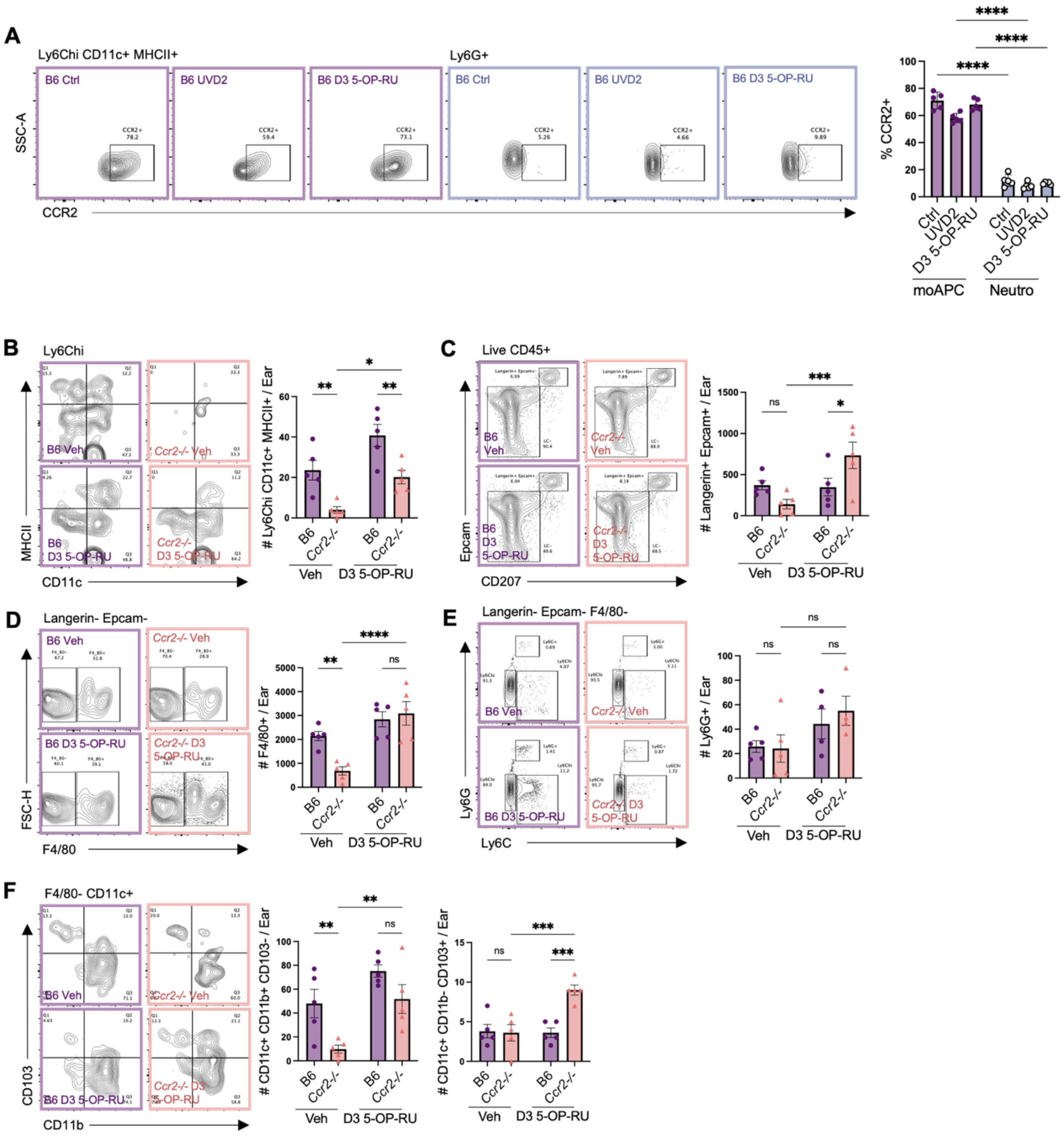

**Figure S17. CCR2 deficiency does not affect 5-OP-RU-driven expansion of non-moAPC myeloid cells.** (**A**) Flow cytometry contour plots and corresponding quantification of CCR2 expression on skin moAPCs and neutrophils at baseline (Ctrl) and early after MAIT cell stimulation (UVD2 or D3 5-OP-RU; n=5/group). (**B-F**) Representative flow cytometry contour plots and corresponding quantification of skin myeloid cells in B6 and *Ccr2-/-* skin early after (D3 5-OP-RU) or without MAIT cell activation (Veh): (B) moAPC, (C) Langerhans cells, (D) macrophages, (E) neutrophils, (F) cDC2 and cDC1 (n=5/group). Data represented as means +/- SEM. Data analyzed with two-way ANOVA. *p<0.05, **p<0.01, ***p<0.001, ****p<0.0001. ns, not significant.

**Table 1. Murine skin Treg immunoregulatory score genes.**

| Immunoregulatory Score | |
|---|---|
| Genes | *Foxp3* |
| | *Il2ra* |
| | *Ctla4* |
| | *Foxo1* |
| | *Tgfbr1* |
| | *Il10* |
| | *Entpd1* |
| | *Tigit* |
| | *Lgals1* |
| | *Tnfrsf1b* |

**Table 2. Human skin MAIT cell and Treg gene signatures.**

| MAIT Cell Signature | | Treg Signature | |
|---|---|---|---|
| Genes | *KLRB1* | Genes | *ACTA2* |
| | *IL18RAP* | | *BCL2L11* |
| | *ZBTB16* | | *CCDC141* |
| | *SLC4A10* | | *CCNG2* |
| | *DPP4* | | *CDK14* |
| | | | *CTLA4* |
| | | | *F5* |
| | | | *FCRL3* |
| | | | *FOXP3* |
| | | | *GBP5* |
| | | | *GCNT4* |
| | | | *GPA33* |
| | | | *ID3* |
| | | | *IKZF2* |
| | | | *IL2RA* |
| | | | *INPP5F* |
| | | | *METTL7A* |
| | | | *OAS1* |
| | | | *PYHIN1* |
| | | | *RBMS3* |
| | | | *RTKN2* |
| | | | *SELP* |
| | | | *SGPP2* |
| | | | *SLC12A6* |
| | | | *SMPD3* |
| | | | *ST8SIA6* |
| | | | *TIGIT* |
| | | | *TNFRSF1B* |
| | | | *TRIB1* |
| | | | *TSHR* |
| | | | *VAV3* |
| | | | *ZCCHC14* |

**Table 3. Mouse qPCR primer sequences.**

| Gene | Forward Sequence | Reverse Sequence |
|---|---|---|
| *Ifng* | CAGCAACAGCAAGGCGAAAAAGG | TTTCCGCTTCCTGAGGCTGGAT |
| *Gzmb* | CAGGAGAAGACCCAGCAAGTCA | CTCACAGCTCTAGTCCTCTTGG |
| *Il6* | TCTATACCACTTCACAAGTCGGA | GAATTGCCATTGCACAACTCTTT |
| *Tnfa* | CTGAACTTCGGGGTGATCGG | GGCTTGTCACTCGAATTTTGAGA |
| *Irf7* | GTCTCGGCTTGTGCTTGTCT | CCAGGTCCATGAGGAAGTGT |
| *Ifi27l2a* | CTGTTTGGCTCTGCCATAGGAG | CCTAGGATGGCATTTGTTGATGTGG |
| *Hif1a* | CCTGCACTGAATCAAGAGGTTGC | CCATCAGAAGGACTTGCTGGCT |
| *Tgfb1* | TGATACGCCTGAGTGGCTGTCT | CACAAGAGCAGTGAGCGCTGAA |
| *Gapdh* | TGGAAAGCTGTGGCGTGAT | TGCTTCACCACCTTCTTGAT |
